# Frequent Whole-Genome Duplication Events Drive the Genomic Evolution of Triple-Negative Breast Cancer During Neoadjuvant Chemotherapy

**DOI:** 10.1101/2025.11.17.688693

**Authors:** Devin P Bendixsen, Fiona Semple, Alastair Ironside, Mustafa İsmail Özkaraca, Natalie Wilson, Alison Meynert, Ailith Ewing, Colin A Semple, Olga Oikonomidou

## Abstract

**Background:** Triple-negative breast cancer (TNBC) is associated with poor survival rate and high genomic instability, generating complex tumour genomes. However, the processes that generate this complexity are poorly studied in longitudinal samples. Here, we study the temporal dynamics of TNBC somatic mutations, revealing major transitions in tumour genome evolution, from diagnostic biopsies, through treatment, to cancer remission or recurrence.

**Methods:** Deep whole exome sequencing and CUTseq, a reduced representation whole genome sequencing approach, were performed in parallel, to comprehensively identify short nucleotide variants (SNVs), copy number alterations (CNAs) and aneuploidies. Tumour samples (N=74) from 22 patients were profiled before and after neoadjuvant chemotherapy (NACT), and encompassed spatially diverse samples from multiple primary breast tumours, to allow tracking of the gain and loss of candidate driver variants over time.

**Results:** Genome-wide SNV mutational burden remained stable across disease progression and RCB classes. However, recurrent SNVs were identified in several known TNBC driver genes in response to treatment, with TP53, MICA, CYP2D6, BRCA1, and BRCA2 being frequently altered. The candidate driver variants in these genes frequently exhibited dynamic changes throughout the course of a patient’s treatment, with the original SNVs in pre-treatment samples often lost, while novel variants in the same genes emerged at subsequent time points.

In contrast to the stable genome-wide SNV burdens, dramatic changes in chromosome structure were seen in all tumours, with abundant CNAs and chromosome arm aneuploidies seen in pre-treatment samples, followed by frequent loss of these alterations post-treatment, and their re-emergence at recurrence. Whole genome duplication (WGD) events appear to drive these dynamics, with a higher frequency of pre-treatment WGD seen in patients with the best response to NACT.

**Conclusions:** Comprehensive longitudinal profiling of the TNBC genome demonstrates the complex interplay of SNVs and structural alterations during tumour progression, leading to diverse evolutionary trajectories impacting patient outcomes. Complex mutational patterns encompassing entire chromosomes emerge during progression, with WGD events making major contributions to intra-tumour heterogeneity, and emerging as a potential candidate biomarker of response at both tumour establishment and recurrence.

## Background

Triple-negative breast cancer (TNBC) accounts for 15-20% of breast cancer cases, and is associated with a high rate of relapse and the worst five-year survival rate of all breast cancer subtypes. For many years, targeted therapeutic options for TNBC patients were limited as TNBCs lack expression of the oestrogen receptor (ER), progesterone receptor (PR), and human epidermal growth factor receptor 2 (HER2), which are essential targets for endocrine and biological therapies. However, more recently TNBC treatments have evolved significantly to include the use of immunotherapy, antibody-drug conjugates and PARP inhibitors combined with traditional chemotherapy, thereby increasing the treatment options beyond targeted treagtments.

Collectively, TNBCs are characterised by high levels of genetic instability resulting in complex patterns of copy number alterations (CNAs) and structural rearrangements [1]. Whole genome duplication (WGD) has been shown to be an independent predictor for poor survival in many cancers including estrogen receptor-positive breast cancers [2]. Although WGD has been shown to occur in TNBC [3], the consequences for tumour evolution and patient outcomes are essentially unstudied. Increased tumour infiltrating lymphocytes (TILs), higher PD-L1 (programmed death ligand 1) expression levels and deficiency in homologous recombination (HRD) are more commonly seen in TNBC compared with other breast cancer subtypes [4]. Differences in molecular subtypes, immune composition and mutational landscapes, underlie the heterogeneity of TNBC. Four TNBC subtypes with unique biology and transcriptional diversity have been identified; basal-like (BL), mesenchymal (MES), luminal androgen receptor (LAR) and immunomodulatory (IM) [5–8]. Each subtype varies in immune composition with improved outcomes reported for TNBC patients with increased expression of immune gene signatures and increased TILs [9–11].

For the TNBC patients with known germline *BRCA1/2* mutations (10%), additional treatment with poly ADP ribose polymerase (PARP) inhibitors can be beneficial [12]. Immunotherapy targeting immune checkpoint inhibitors (ICIs) has revolutionised the treatment of TNBC patients, but response is suboptimal in some patients, therefore the search for further biomarkers of response beyond PD-L1 is ongoing [13]. TNBC patients often respond well to neoadjuvant chemotherapy (NACT), with 34% of patients achieving pathological complete response (pCR) which is subsequently associated with 75% lower risk of recurrence, compared to TNBC patients not achieving pCR [14,15]. The recent Keynote-522 trial demonstrated that combining the ICI Pembrolizumab with conventional chemotherapy significantly increased pCR rates with 64.8% of patients acheiving pCR in the experimental arm compared to 51.2 % in the placebo arm. Furthermore, the addition of Pembrolizumab, increased event-free survival rates to 84.5% compared to 76.8% in the control arm, with significant effects on overall survival [16,17]. High pCR rates have been associated with higher tumour mutational burden (TMB), elevated levels of PD-L1 expression and increased levels of TILs in the tumour microenvironment [18]. Defining the landscape of TNBC and the molecular changes that occur in response to NACT, will allow stratification of patients for directed treatment. However, this is confounded by the high genomic instability and extensive intra-tumour heterogeneity (ITH) that are thought to drive the pathogenesis, treatment resistance and metastatic spread of TNBC [19]. When and how these genetic alterations emerge is not known, but WGD is thought to be common in TNBC [3,20], while aneuploidies and copy number alterations (CNAs) have been implicated as drivers of the adaptive response to chemotherapy and metastasis [21,22].

Studies including multiple samples over time have the potential to reveal new tumour biology, but can also entail large investments of time, effort and resources. There is consequently a pressing need for accessible, affordable assays with relatively modest requirements for computationally intensive analysis. For example, although the mutational landscape of TNBC has been examined in previous studies (e.g. Lehmann et al, 2021), the temporal dynamics of CNA and short nucleotide variants (SNVs including single nucleotide substitutions and indels) and WGD during disease progression are poorly studied. Consequently, we lack a detailed account of tumour evolution as patients are diagnosed, undergo NACT, and their disease enters remission or recurs. Here, we employ deep (∼100X) Whole Exome Sequencing (WES) to assay recurrent SNVs and CNAs in patient TNBC samples to identify variants that play a role in disease progression and response to treatment. In parallel, we use CUTseq [23], a reduced representation Whole Genome Sequencing (WGS) approach, to comprehensively identify copy number alterations genome-wide. In addition to recurrent SNVs in known TNBC driver genes we demonstrate major shifts in CNAs throughout the genome following NACT, suggesting new biomarkers of treatment response and revealing critical roles for WGD in tumour evolution.

## Methods

### Patient and Tumour samples

Samples from TNBC patients treated at the Western General Hospital Cancer Centre, enrolled in the NEO and metcfDNA studies (references: 2014/0005 and SR680) at Edinburgh University were acquired through the NRS BioResource and Tissue Governance Unit. Patients with TNBC were determined from pathology records and only samples with matched tumour and blood samples were included. One sample with no matched blood was paired with the patient’s normal breast sample. Collection of human tissues and blood and relevant clinicopathological data were approved by the South East Scotland Research Ethics Committee 01 with REC references: 13/SS/0236 and 15/ES/0094, and all patients consented to sample collection and use of their data in research. Patients and their clinical data (Table S1) were anonymised. All patients underwent NACT with anthracyclines followed by taxanes in combination with Carboplatin, and response was assessed using the internationally recognised residual cancer burden (RCB) scoring index. This system provides a continuous score (RCB score) and categorises patients into four discrete groups (RCB 0, RCB I, RCB II, RCB III) based on the amount of residual disease following NACT. RCB-0 corresponds to no residual disease, also known as pathological complete response (pCR), RCB-1 denotes a small amount of residual invasive cancer, RCB-II a moderate amount of residual invasive cancer and RCB-III indicates substantial residual disease [24]. Samples from before treatment with NACT (PreT), at the end of NACT after surgery (PostT), where available from recurrent tumours (R) and from selected mid NACT biopsies (at completion of anthacyclines before commencing taxanes/carboplatin) (MidT) were collected. Tumour infiltrating lymphocytes (TILs) were assessed by a breast cancer consultant pathologist (AI) using light microscopy on routine diagnostic H&E-stained glass slides. The number of TILs were quantified as a percentage of the tumour-associated stroma occupied by lymphocytes, as per the recommendations of The International Immuno-Oncology Biomarker Working Group on Breast Cancer. Three distinct regions of predefined size for each tumour were evaluated for TILs using a visual reference image index of pre-determined densities [25,26].

### DNA extraction

Formalin fixed paraffin embedded (FFPE) samples were sectioned onto slides using a cryostat. An initial 4µm section was stained with hematoxylin and eosin (H&E) and reviewed by a breast cancer consultant pathologist (AI) to determine sample tumour cellularity and mark up the tumour regions (Table S2). DNA was extracted (using Qiagen QIAamp DNA FFPE Advanced UNG Kit #56704) from 10µm sections, 2 to 5 sections per sample depending on tumour cellularity. Extraction was carried out on marked up tumour enriched areas and where possible, multiregional sampling was carried out by extracting DNA from spatially distinct tumour enriched areas on each slide, or from slides prepared from different FFPE blocks from the same tumour sample. Germline DNA was extracted from whole blood collected before commencement of NAC using the Qiagen DNeasy Blood & Tissue Kit (#69504). DNA was quantified using the Qubit dsDNA HS Assay Kit (#Q32851).

### Whole Exome Sequencing

The quality of the DNA extracted from the FFPE samples was assessed with the Illumina FFPE QC Kit (Illumina Inc, #15013664) according to the manufacturer’s protocol. The required ΔCq ≤5 was achieved by all samples determining suitability for library preparation. For each sample, 50ng of DNA sample was used for library preparation with the Illumina DNA Prep with Enrichment, (S) Tagmentation Kit (Illumina #20025524) using IDT® for Illumina® DNA/RNA UD Indexes Set A, Tagmentation (Illumina #20027213) according to the manufacturer’s protocol with modifications for FFPE DNA where appropriate. Using the Twist Comprehensive Exome panel (Twist Biosciences #102032), 500ng of each pre-enrichment gDNA library was pooled in sets of 12 and hybridised to the targeted regions during an overnight incubation, followed by capture onto magnetic beads and amplification of the exome enriched library pools, which were then purified with Agencourt AMPure XP beads (Beckman Coulter, #A36881). Libraries were quantified using Qubit dsDNA HS assay and quality assessed using Agilent Bioanalysis with the DNA HS Kit (#5067-4626). Fragment size and quantity measurements were used to calculate molarity for each enrichment library pool. Sequencing was performed on the NextSeq 2000 platform (Illumina Inc, #20038897) using NextSeq 2000 P3 Reagents (200 Cycles) (#20040560). A single pool containing the 8 enrichment pools was sequenced on 3 separate P3 flow cells to achieve a minimum of 100X coverage. PhiX Control v3 (Illumina, # FC-110-3001) was spiked into each run at a concentration of 1%. Library preparation and sequencing were carried out at the Clinical Research Facility (CRF), Edinburgh, UK.

### Reduced Representation Whole Genome Sequencing – CUTseq

Multiplexed DNA sequencing libraries were generated using CUTseq [23] following the published protocol (https://protocolexchange.researchsquare.com/article/pex-25/v3) with several modifications. Firstly, adapters (A1-A48) specific to the HindIII cut site which incorporate sample-specific barcodes, the RA5 Illumina sequencing adapter and the T7 promoter sequence were purchased from Integrated DNA Technologies (IDT) (Table S3). Briefly, forward strand barcode adapters were 5’ phosphorylated then annealed to the reverse strand barcode adapters by incubation at 95°C with step down cooling to 25 °C over the course of 45min to form double-stranded(ds)-barcoded HindIII-specific adapters. DNA (10-50ng) from patient samples was fragmented by enzymatic digestion with HindIII-HF® (NEB, # R3104L) for 4hr at 37 °C in a 10μl volume of CutSmart® Buffer (NEB, #B6004). Generally, 48 samples were prepared at the same time. Digested DNA samples were barcoded using T4 DNA ligase (NEB, # M0202T) by incubating with ds-barcoded HindIII adapters at 16°C for 18hr. Samples were combined into 3 pools of 16 samples, each containing ∼300-500ng starting DNA, and purified using Agencourt AMPure XP (Beckman Coulter, #A63880) at a ratio of 0.9X beads to sample. The 3 pools were combined and fragmented with target peak size 200bp using the Covaris E220 Evolution. Sample concentration was determined using Agilent Technologies DNA HS Kit (#5067-4626), followed by evaporation to ∼20ng/μl. In vitro transcription was carried out with 8μl of sample (∼160ng) using the MEGAscript® T7 Transcription Kit (ThermoFisher Scientific, #AM1334-5) following the manufacturer’s protocol, and the resulting RNA was purified using Agencourt RNAClean XP beads (Beckman Coulter, #A63987) at a ratio of 1.8X beads to sample. For RNA sequencing library preparation, primers (RTP, RP1 and RPIndex) and adapter (RA3) were purchased as standard desalted oligos from IDT (Table S3). RNA was quantified using the NanoDrop Spectrophotometer (Thermo Scientific) subsequently 1μg of RNA was used for library prep. Briefly, the RA3 adapter was ligated using T4 RNA ligase 2, deletion mutant (Cambio Ltd, #LR2D1132K) at 25°C for 2hr, followed by reverse transcription using the RTP primer and SuperScript IV RT (Invitrogen, #18090050) at 50°C for 1hr. Each library was then amplified using NEBNext® UltraTM II PCR master mix (#M0544L) and indexed using Illumina primers (RPI2 and RPI3) to allow simultaneous sequencing of two 48 sample libraries. After double-sided size selection purification using AMPure XP beads, final library quantification and size distribution were carried out by Qubit dsDNA HS assay and Agilent Bioanalysis (DNA HS Kit, #5067-4626). Sequencing (1×131bp) was performed on the NextSeq 2000 platform (Illumina Inc, #SY-415-1002) at the CRF, Edinburgh, using the NextSeq 1000/2000 P3 Reagents (100 cycles) v3 Kit (#20040599) with PhiX Control v3 (Illumina, #FC-110-3001) spiked in at a concentration of 5%.

### Primary processing of WES and CUTseq sequencing data

The raw sequencing reads for the WES and CUTseq datasets were assessed and processed independently. Sequencing reads generated by CUTseq were pre-processed using an established pipeline (<u>github.com/ljwharbers/copynumber-calling</u>). Briefly, reads were demultiplexed into individual samples based on the internal CUTseq 8 bp barcodes allowing for a single mismatch. The adapters were trimmed using fastp v0.23.2 [27] and reads were aligned to the GRCh38 reference genome using BWA mem v0.7.17 [28] and SAMtools v1.16.1 [29]. Duplicate marking and mapping and alignment quality were assessed using Picard and Alfred v0.2.6 [30]. The resulting mapped reads provided a genome-wide readout (at 3,422bp resolution) across all HindIII restriction enzyme cut sites.

The raw sequencing reads from WES were processed using the nf-core [31] pipeline Sarek v3.2 for identification of germline and somatic variants [32,33]. Reads were assessed for quality using FastQC v0.11.9 [34] and MultiQC v1.17 [35] and adapter and quality-trimmed using fastp v0.23.2 [27] then aligned to the GRCh38 reference genome using BWA mem v0.7.17 [28] and SAMtools v1.16.1 [29]. Base recalibration and duplicate marking was performed using GATK4 v4.3.0.0 [36] and exome-wide read depth was assessed using mosdepth v0.3.6 [37].

### Variant calling and filtering

To assess the mutational landscape within our samples, SNVs and small insertions and deletions (indels) were identified from the WES samples. To limit bias in the identification of somatic variants and to rescue potential false negative mutations, SNVs and indels were called as a majority vote between Mutect2 (GATK v4.3.0.0) [36,38], Strelka2 v2.9.10 [39] and VarScan v2.4.6 [40]. Somatic variants were annotated using Ensembl Variant Effect Predictor (VEP) v107 [41]. CaMutQC v1.1.0 [42] was used to filter somatic variants based on sequencing quality thresholds, strand bias or the presence of an adjacent indel tag, applying the following parameters: tumour total depth ≥ 40, tumour variant depth ≥ 5, normal total depth ≥ 10, Variant Allele Frequency (VAF) ≥ 0.1 and strand bias score ≥ 2. Variants occurring in the 1000 Genomes Project dataset [41] were removed as these were highly likely to be germline. To be retained, a consensus somatic variant (identified by ≥ 2/3 variant callers) had to be in a protein coding region, pass CaMutQC filtering thresholds, have an allele frequency ≥ 0.1, and be estimated by VEP to have high or moderate impact. For each patient, germline variants were identified from matched blood or normal tissue samples using VarScan v2.4.6 [40] and annotated using Ensembl VEP v107 [41] then checked against the ClinVar database v20240107 to identify clinically relevant mutational variants [43]. Tumour mutational burden was calculated as the number of consensus mutations per Mb and microsatellite instability (MSI) was assessed using MSIsensor-pro v1.3.0 [44].

Genes identified as a source of artefacts [45–47] were removed, and mutational loads in TNBC relevant genes [7,8,48–51] were assessed in detail (Table S4). The number of alterations in oncogenic signalling pathways was determined using Maftools and OncogenicPathways [52,53]. SNV mutational signatures were extracted using the mutSignatures pipeline (v2.1.5) [54]. Preliminary mutational process assessment suggested that two mutational signatures reconstruct the SNV landscape with minimal error (compared to median mutation counts) and was further corroborated by a silhouette plot after extraction. The de novo signatures were then compared to the COSMIC SBS Mutational Signatures v3.4 [55,56].

To infer candidate driver variants two driver prediction methods with different approaches were applied, dNdScv [57] and OncodriveFML [58], on SNVs with variant allele frequency (VAF)>0.1 in PreT samples. OncodriveFML v2.2.0 was used with default parameters to identify candidate variants based on significant bias in functional impact of mutations (Q-value<0.05). dNdScv was used with default parameters and identified candidate driver variants based on the ratio of synonymous to non-synonymous mutations (qglobal_cv<0.05).

To detect structural variants, Manta v1.6.0 [59] was used on the WES samples. Five categories of structural variants were assessed: inversions (INV), insertions (INS), deletions (DEL), duplications (DUP), and breakend (BND) variants. To limit the number of false positives, ‘micro-inversions’ less than 400bp were removed from the analysis. Each WES and CUTseq sample was assessed for somatic CNA using ASCAT.sc v0.1 [60]. This approach can be employed on either shallow-coverage or targeted sequencing and uses average read depth over various bin sizes as opposed to per base pair read depth. Crosetto et al have shown that CUTseq with HindIII digestion yields reproducible copy number profiles at 100Kb resolution but we conservatively adopted a resolution of 300Kb guaranteeing a larger number of CUTseq sites in each bin (the average distance between 2 consecutive HindIII sites is 3.4kb) [23]. CNA calls based upon WES or CUTseq were removed if they overlapped a blacklisted region [61], known to produce mapping artefacts, by >25% of the CNA length. The union of CNA calls from WES and CUTseq was used for all subsequent analyses. CNA calls were intersected with a list of genes (Table S9) implicated in TNBC progression compiled from the literature [8,49,62]. CNA deletion/amplification hotspots were determined by GISTIC v2.0.23 [63] based on WES and CUTseq data for PreT samples. Regions were considered significantly enriched for alterations at a q-value < 0.05 [63].

Homologous recombination deficiency (HRD) was predicted using scarHRD v0.1.1 [64] based on loss of heterozygosity (LOH), large scale transitions (LST) and telomeric allelic imbalances. WES sequencing copy number profiles from Sequenza [65] were used as input to scarHRD. WGD events were called where ASCAT.sc estimated ploidy for either WES or CUTseq were ≥3, which is a more conservative approach than a previous TNBC study using >2.7 [7].

### Analysis of co-occurrence or mutual exclusivity

Linear relationships among genomic metrics (i.e. TMB, CNA burden, ploidy, etc) and clinical variables (TIL, RCB, OS), were determined using the R package ‘corrplot’ [66] using a confidence level of 0.95, adjusted for multiple testing using the Benjamini-Hochberg procedure. Patterns of pairwise co-occurrence or mutual exclusivity were tested using SELECT v1.6.3 [67,68]. SELECT was run 10 times using 10 different random seeds and the median select score for each pairwise interaction was used. The effects of variables on overall survival were modelled with Cox proportional hazards regression models using the coxph method in the R package ‘survival’ v3.8.3 [69,70] and Kaplan-Meier curves were plotted using R package ‘ggsurvfit’ v1.1.0 [71].

### Analysis of mutational patterns across time

We employed two strategies to examine mutational patterns over time, one based upon differences between mean values at each timepoint, and another considering entire temporal trajectories for each patient, based on linear mixed models (LMM). In the first strategy, for each genomic metric (i.e. TMB, CNA burden, ploidy, etc), significant differences between sample means at each timepoint (PreT, PostT, R) or across RCB classes were assessed using Kruskal-Wallis rank sum tests. In cases of presence/absence (i.e. SNV prevalence or copy number status) a Fisher’s exact test was used for two variables and a Chi-squared test for three or more. Pairwise comparisons amongst groups were made using Wilcoxon rank sum tests with Bonferonni corrections for multiple testing. In the second strategy we applied LMM (for continuous variables) or Generalised Linear Mixed Models (GLMM) (for binary dependent variables), treating patients as random effects, to examine complete temporal trajectories. All linear mixed models were implemented in Python using the mixedlm function from the statsmodels package [72], which supports the random-effects structure required for repeated measures data. For generalised linear mixed models, we used the pymer4 package [73], which interfaces with the R lme4 framework [74] to support non-Gaussian outcomes such as binary variables. This approach allowed us to accurately evaluate whether any of the molecular features changed significantly over time, taking the entire temporal trend into account for each patient, and adjusting for within patient correlations between timepoints. The variables analysed and the corresponding summary results from the LMM/GLMM models are provided in Table S14.

## Results

Tumour samples were collected from 22 TNBC patients over the course of NACT and progression of the disease. Samples collected at the following timepoints, PreT (before NACT), PostT (end of NACT after surgery), R (from recurrent tumours) and MidT (at the midpoint of NACT) are shown in **Figure 1A**. Response to NACT was assessed by a breast cancer consultant pathologist and the residual cancer burden (RCB) score and RCB class were determined for each patient (Table S1). Patients were ranked from low to high RCB (**Figure 1B** and Table S1). PreT samples were available for all patients while PostT samples only exist for patients where there was not pathological complete response to NACT (RCB=I-III). Where available, multiple regions (n=2-4) of the PostT specimen were obtained and analysed. Ten patients had recurrence following completion of NACT and subsequent adjuvant treatment. Recurrence samples (R) were attainable for 7 of these patients, and it was possible to sample multiple regions (n=2-4) in 3 patients (Table S1). The aim was to establish the most complete sampling over time for each patient, supported by multi-region sampling at timepoints where this was feasible. The numbers of samples sequenced and analysed for each patient at each time point are shown in **Figure 1B** and the full clinical and pathological characteristics of TNBC patients in the study is described in Table S1. The level of tumour infiltrating lymphocytes (TILs) was measured in all samples and varied widely, from 1-90% of the stromal area (**Figure 1C**, Table S2). Highest infiltration was observed in PreT samples in patients with a good response to NACT and was negatively correlated with RCB score as expected (Figure S1). Accordingly, tumours recurred in 4/5 patients with PreT TIL levels of <10%, consistent with previous reports [75] and increased TIL levels in PostT in our study was not indicative of recurrence as previously reported [76]. In the patients with recurrence samples, TILs significantly decreased from a PreT average of 33% to an R average of 5% (Figure S1, p=0.03).

**Figure 1.**
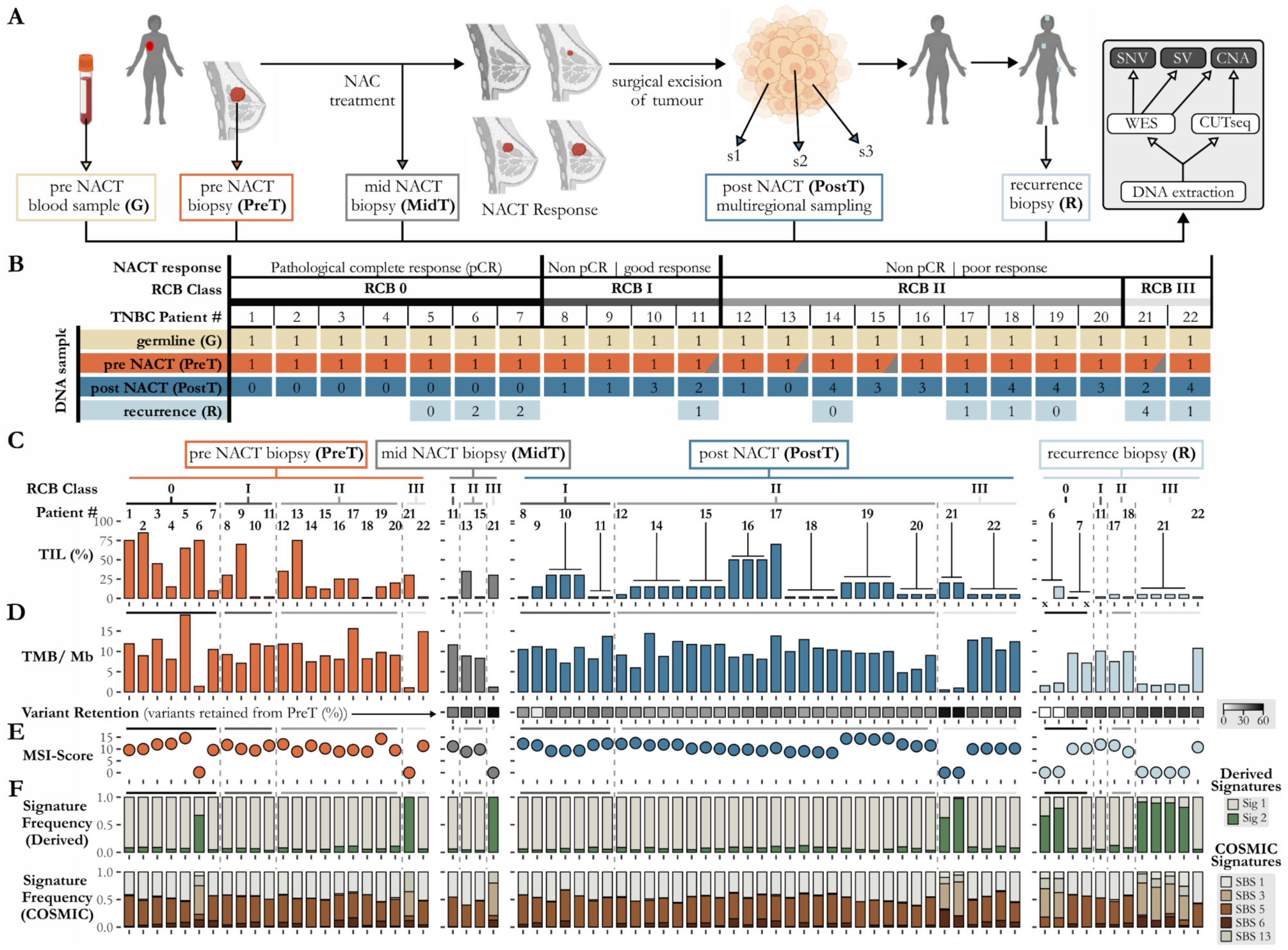
Study overview and tumour mutational burden in triple-negative breast cancer. **(A)** Longitudinal samples were collected over several years for each patient, including a blood sample for germline DNA, a PreT biopsy before commencement of NACT and a MidT biopsy (in 4 patients). Multiple regions (n=1-4) of the PostT and R tumour samples were sequenced. **(B)** Patients (n=22) were selected based on response to NACT; good response = residual cancer burden score of RCB 0/I, poor response = RCB II/III. Samples (n=96) are represented by coloured rectangles, yellow = germline, orange = PreT, orange with grey triangle =MidT, blue = PostT, light blue = R. The number of regions sampled per timepoint per patient are indicated in the rectangles (n=0 means no sample was available due to complete response to therapy or unattainability at recurrence sites). **(C)** The percentage (0-100%) TILs found in each tumour sample, at each timepoint, orange = PreT, grey = MidT, blue = PostT, light blue = R, is shown for each patient with TNBC patient number indicated at the top. A patient number on top of a T-bar indicates where multiregional samples were assayed from that patient. Vertical dotted lines within each timepoint separate RCB classes. **D)** TMB calculated as SNVs per Mb for each patient across timepoints, orange = PreT, grey = MidT, blue = PostT, light blue = R. The proportion of SNVs retained from PreT across subsequent timepoints is shown. Increasing variant retention percentages are indicated on a scale of white to black (0-60%). **(E)** MSI scores for each sample across the time course, dots representing scores between 0 and 15 are coloured, as above to indicate different timepoints. **(F)** Mutational signatures found using *de novo* extraction (top) or by estimating the contribution of known COSMIC SBS mutational signatures (bottom).

### SNV mutational burden remains stable between RCB classes and across disease progression

Tumour mutational burden (TMB) was assessed for each tumour sample (n=74) using the total burden of consensus SNV (short nucleotide variant) calls from deep whole exome sequencing (WES) (**Figure 1D**, Table S5, Methods). TMB for PreT samples ranged from 1.1 to 19 SNV/Mb, with most patients ranging from 7.1 to 15.6 SNV/Mb. As might be expected, TMB and patient age at diagnosis were positively correlated (R^2^=0.22, p=0.027). Two notable outlier patients exhibited significantly lower TMB (1.1 and 1.4 SNV/Mb) and were found to harbour germline mutations in *BRCA1* and *RAD51C* (TNBC6 and TNBC21 respectively) predicted to cause homologous recombination repair deficiency (HRD) (Figure S2). In general, across each patient’s longitudinal samples the mean TMB did not vary significantly across the course of the disease, based upon pairwise comparisons between timepoints (**Figure 1D**); though analysis of complete longitudinal profiles using linear mixed modelling (LMM) revealed a modest decline in TMB for R samples (p=0.03) compared to PreT samples (Table S14). Overall, TMB did not differ significantly between different RCB classes (Figure S3). Mean genomic SV burden also showed no significant relationship with sampling time (PreT, MidT, PostT or R) or RCB class (Figure S4, Table S8). Microsatellite instability (MSI) was largely stable across time (**Figure 1E**, Table S5), with slight decreases in PostT (p=0.026) and R samples (p=0.018) compared to PreT detected in longitudinal LLM modelling (Table S14).

We exploited our deep WES data to study high confidence SNVs (supported by >40 reads) in detail, providing a window on variants lost or newly emerging during disease progression. The fate of SNVs (single nucleotide substitutions and indels) in each patient was followed using a novel metric: variant retention (**Figure 1D** and Table S5). Variant retention is defined as the proportion of SNVs found in common between PreT and subsequent timepoints from the same patient. Between PreT and PostT the retention of SNVs in the majority of patient PostT samples was generally low, ranging from 5-56%, suggesting frequent replacements of one dominant clone by another within tumours in response to treatment. In patients with both PostT and R samples the variant retention at recurrence generally approximated PostT values (on average ±1.12%). Additional samples midway through NACT (MidT) in a fraction of patients (n=4) allowed an insight into how quickly variants can be altered during treatment. The variant retention for the MidT samples were equivalent or higher than paired PostT samples for those patients, consistent with rapid responses to treatment.

Mutational signatures were extracted *de novo* from the SNV mutational data (Methods), and the prevalence of known COSMIC SBS mutational signatures was also assessed (**Figure 1F**, Table S7). Derived signature 1 (Sig 1) closely matched COSMIC SBS1 (0.89 cosine similarity) and derived signature 2 (Sig 2) most closely matched SBS13 (0.70 cosine similarity). SBS1 and SBS5 signatures were prevalent in most patients, with minor influences by SBS3, SBS6 and SBS13. SBS1 and SBS5 have been shown to be clock-like signatures and increase with patient age. In patients harbouring germline disruptive SNVs in BRCA1 and RAD51C (TNBC6 and TNBC21), SBS3 was the most dominant signature, which is associated with defective homologous recombination repair. Overall, the stable SNV mutational burden over the progression of TNBC was matched by consistent mutational signatures, suggesting that the driver SNVs present in each tumour may also show stability over time.

### Loss and emergence of SNVs in known driver genes is common during TNBC treatment

Analysis of SNVs disrupting protein coding regions revealed many candidate driver genes found in previous studies. After filtering to limit false positive SNV calls (Methods), we identified SNVs disrupting 57 of 98 candidate genes previously associated with TNBC progression in the literature (Figure S2, Table S5). Those candidates disrupted most frequently by SNVs found in the 22 patients here are shown in **Figure 2A**. A combination of driver gene prediction algorithms (Methods) identified *TP53*, *CYP2D6* and a novel TNBC-associated gene *MICA* as candidate driver genes (Table S6), in spite of the limited statistical power of this modest cohort. Variants in one or more of these 3 genes were observed in 17/22 patients. The most frequent variants found in the PreT samples were in *TP53* and *MICA*, both genes being altered in 9/22 patients (41%), followed by *CYP2D6* with variants in 7/22 patients (32%). *BRCA1* and *BRCA2* variants were mutually exclusive (Figure S5), with *BRCA1* or *BRCA2* being detected in 10/22 patients (45%). Pairwise analysis in the PreT samples determined disruptive *BRCA2* variants (in 6 patients) and disruptive *TP53* variants (in 9 patients) to also be mutually exclusive events (Figure S5). Overall, 91% (20/22) of patients had PreT samples with somatic or germline disruptions of *TP53*, *BRCA1*, *BRCA2*, *PRKDC* or ATM, consistent with the early development of genomic instability. The remaining two patients (TNBC9, TNBC15) developed disruptive variants in these genes at PostT, suggesting ongoing selection for genomic instability via alterations to these same 5 genes across the cohort.

**Figure 2.**
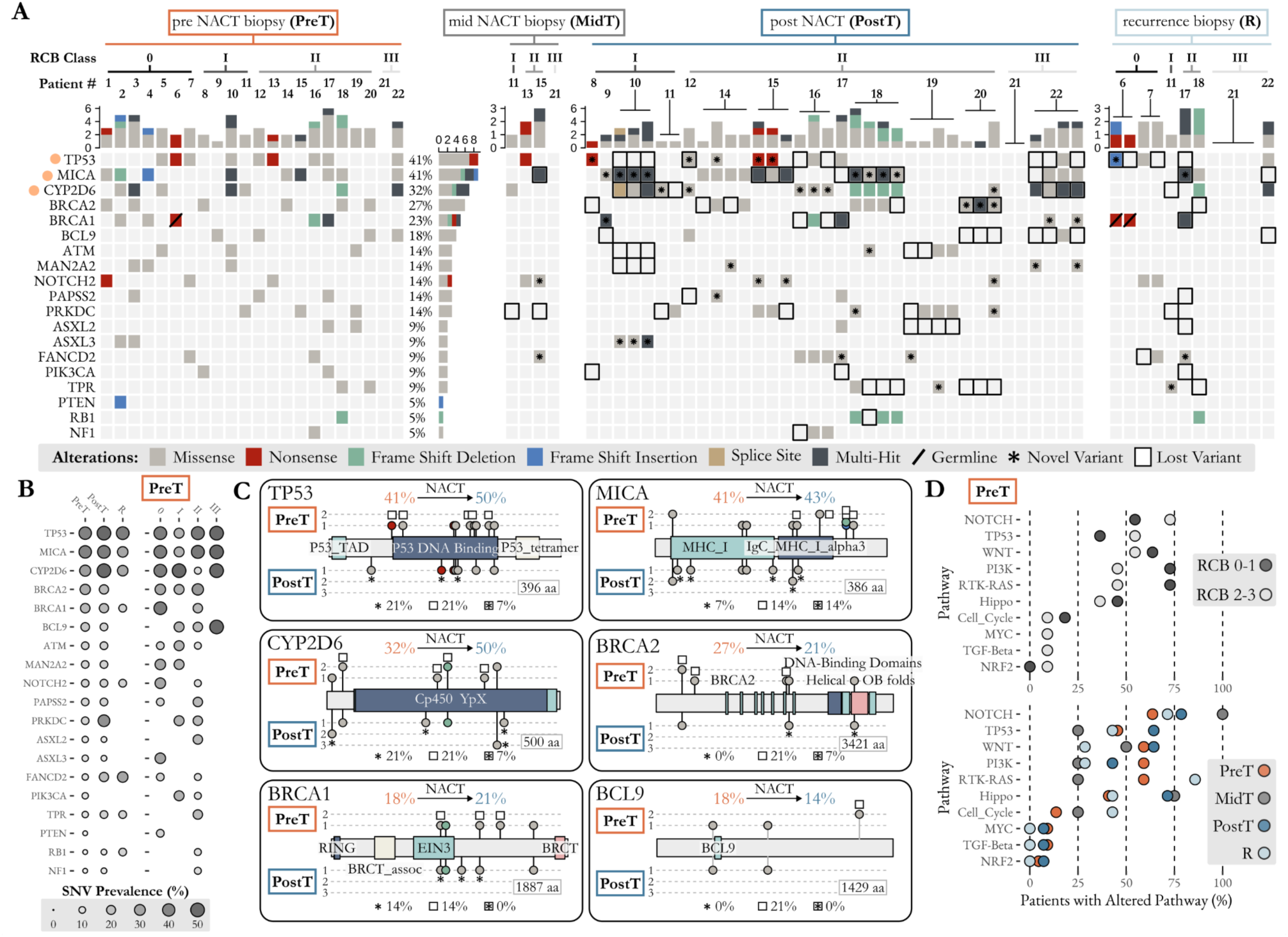
Longitudinal SNV landscape of TNBC during NACT. **(A)** Oncoplot indicating the presence of deleterious variants predicted to disrupt TNBC gene function. Variants detected for each TNBC patient (patient number indicated at the top) are shown at each timepoint, with variant type indicated by a coloured square (see key, bottom). For PostT and R samples a patient number on top of a T-bar shows the number of multiregional samples assayed from that patient. Variants lost between PreT and PostT or between PreT and R, are denoted by a square with a black outline. Variants not found in PreT that are detected in PostT or R, are denoted by an asterisk. Orange dots indicate genes identified as driver candidates and germline variants are indicated with a solid diagonal line. Where multi-region samples exist for the same timepoint they generally support the existence of the same driver mutation. **(B)** SNV prevalence through longitudinal time (PreT, PostT, R) and across RCB classes (0, I, II, III). **(C)** Gene diagrams of the six most commonly altered genes comparing the location and prevalence of mutations in the PreT and PostT samples. Variants type is indicated and coloured as in panel A. White squares indicate lost variants and asterisks indicate newly emerging variants occurring between PreT and PostT timepoints. Orange and blue numbers indicate the proportion of patients with altered genes in PreT and PostT respectively. Percentages beneath each diagram indicate the proportion of PostT patients with ≥1 sample with a novel (asterisk), or lost (square) variant, as well as variants observed to be both novel and lost in the cohort (asterisk in square). **(D)** Proportion of altered oncogenic pathways across RCB classes in PreT samples (top) and through time (bottom).

We employed a conservative consensus SNV calling (sequencing read depth >40, variant allele frequency (VAF)>0.1) strategy to ensure confidence in the detection of both ‘lost’ and ‘novel’ variants in later (PostT and/or R) samples, relative to the initial PreT biopsy samples. Losses and gains of SNVs were frequently observed over time, even in the 5 most frequently altered driver genes TP53, MICA, CYP2D6, BRCA1 and BRCA2 (**Figure 2A**). For example, TP53 variants detected in the PreT samples of 7 patients were no longer observed in the paired PostT or R samples of these patients. This included a patient (TNBC10), where three PostT samples were sequenced and the TP53 variant identified in the PreT sample of the patient was lost in all three samples. Whereas, in patient TNBC22, four PostT samples were sequenced and the corresponding PreT TP53 variant was only identified in one sample suggesting intra-tumour heterogeneity. In addition, TP53 variants arose in the PostT or R samples of 5 patients where these variants were not detected in the paired PreT samples. Remarkably, in some patients, variants were lost concurrently with new variants arising in the same gene. This was seen with MICA SNVs in 4 patients, where later (PostT or R) gains and losses of variants were observed relative to the paired PreT samples. Some genes, i.e. BCL9 and PIKC3A were subject only to loss of the PreT variants across the time course of the study, with 3/4 patients’ (BCL9) and 2/2 patients’ (PIK3CA) variants undetectable after NACT. PostT and R samples also gained variants in a number of TNBC-related genes, in particular ERBB2 variants appeared in 4 patients, ALK variants in 3 patients and EGFR variants in 2 patients, with additional genes obtaining PostT variants in single patients (Figure S2). Thus, the consistent SNV burdens seen genome-wide (**Figure 1**) obscure frequent turnover at the level of particular driver variants in particular genes, likely reflecting clonal competition and replacement within tumours over time.

The prevalence of genes altered by SNVs were largely consistent over time and across RCB classes (**Figure 2B**). A notable deviation is the prevalence of BRCA1 variants (3/7 patients) in the PreT samples of patients in RCB 0, which exhibited the best response to NACT. The locations of the SNVs occurring within each of the 6 most altered driver genes show similar patterns over time (**Figure 2C**, and Table S5). In general, it seems that both lost and novel SNVs disrupt similar regions of the proteins encoded by these driver genes, and this is also true of SNVs maintained across all timepoints in the same genes. For example, across both PreT and PostT samples, 10 different TP53 variants, including 8 missense and 2 nonsense variants, were detected. Multiple lost and novel SNVs were found, and on 4 occasions the same variant was observed in both PreT and PostT, but regardless of timing the vast majority were present in the DNA binding domain (**Figure 2C**). Other driver genes failed to show strong biases in the locations of lost and novel variants, suggesting similar disruptions of gene functions over time. There were no statistically significant associations between candidate driver variants and patient survival, though such analyses are under-powered with the current data (Figure S6). Shifts in altered oncogenic pathways were also evident, across time and across RCB classes, based upon changes in the proteins predicted to be altered by SNVs (**Figure 2D**). Patients with poorer responses to NACT (RCB 2-3) show more frequent alteration of the NOTCH (+18%, OR=1.3) and TP53 (+18%, OR=1.5) pathways than patients with better response to NACT (RCB 0-1). Better responding patients (RCB 0-1) showed higher frequencies of altered WNT (+9%, OR=1.2), PI3K (+27%, OR=1.6) and RTK-RAS (+27%, OR=1.6) pathways (**Figure 2D**).

### Comprehensive detection of copy number alterations over disease progression

Copy number alterations (CNAs) were detected in our cohort using two different but complementary assays, WES and CUTSeq [23] (**Figure 3A**). WES is known to provide a detailed interrogation of mutations within the fraction of the genome occupied by exons, while CUTSeq provides a broader picture of CNAs that are distributed throughout the human genome. For example, a closer examination of chromosome 8 in a patient, revealed that both approaches identified an amplification of chromosome arm 8q (**Figure 3B**). However, as expected the read depth of WES closely mirrors the density of genes, whereas CUTseq shows a more uniform distribution across the chromosome arm. Decreasing the genomic range and examining a more focal region around a single gene (*MYC*) further exemplifies the high concentration of reads within exons in WES, while CUTseq shows a more even distribution. In contrast, a closer examination of chromosome X revealed a deletion near the centromere that was detected using CUTseq, but not by WES (**Figure 3B**). The detection of such alterations in gene-poor regions highlights the potential of CUTseq to assess genomic instability more comprehensively.

**Figure 3.**
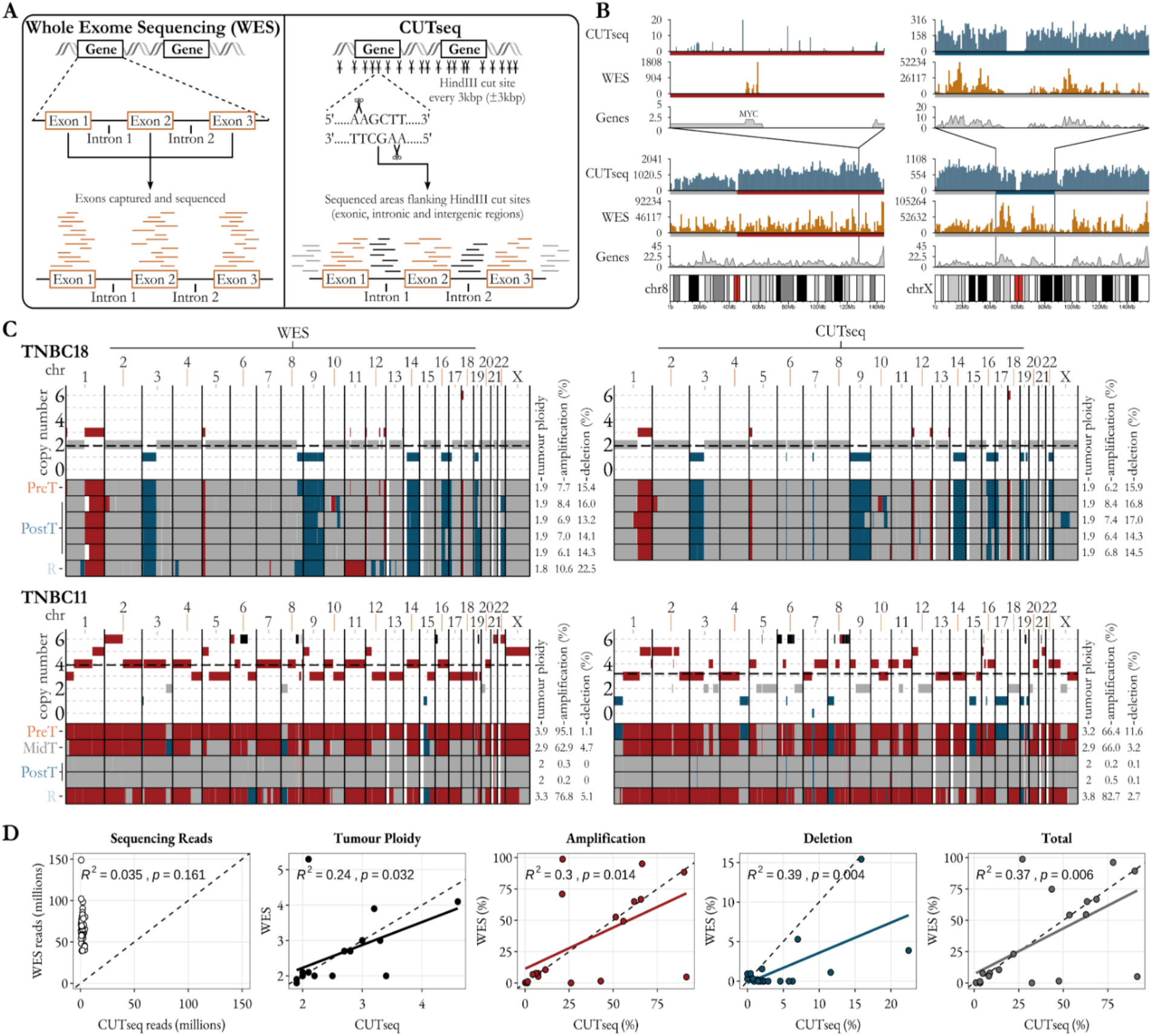
Comparison of whole exome sequencing (WES) and CUTseq in detecting CNAs. **(A)** Graphical representation of WES and CUTseq approaches. **(B)** Examples of sequencing read depth and coverage of chr8 and chrX for a single patient. Chromosome karyotypes and gene density are shown for context. **(C)** Longitudinal CNA calls for two patients (TNBC18 and TNBC11) indicating the consistent patterns exhibited between WES and CUTseq. Red indicates amplified genomics regions, blue indicates deleted regions, grey indicates diploid regions, black indicates regions with estimated copy number >6, and white indicates regions where copy number was unobtainable. The dashed black line indicates the estimated tumour ploidy. The 4 plots show the estimated copy number (y-axis) for each chromosome (x-axis) for the PreT sample of the 2 selected patients. Below this, amplifications and deletions are summarised for all timepoints, including multiregional samples, as indicated on the left. Estimated tumour ploidy, and the percentage of the genome amplified or deleted from each approach is shown on the right. **(D)** Pairwise sample comparisons of CUTseq and WES for number of sequencing reads, estimated tumour ploidy, percentage of genome amplified, deleted and the total altered by CNAs.

CNA patterns over the course of treatment showed differences between patients (all CNA profiles are provided in Figure S7). Patient TNBC18 (**Figure 3C**) provides an archetypal example of stability over time, where amplified and deleted regions present at PreT, were also found in all 4 tumour regions sampled at PostT, and at recurrence (with similar profiles over time seen in both CUTSeq and WES). In contrast, CNA profiles for many patients, such as TNBC11, were dynamic over time (**Figure 3C**), with widespread amplifications seen across most chromosomes in the PreT sample becoming completely absent in PostT samples, then re-appearing in the R tumour sample. Importantly, this pattern of alteration loss and recurrence through longitudinal time is consistent in both CUTseq and WES data for TNBC11, demonstrating broadly consistent results between both assays. Across all samples, there was a high level of agreement between the two approaches (**Figure 3D**), as indicated with significant correlations of estimated tumour ploidy (p=0.032) and the proportion of each genome either amplified (p=0.014) or deleted (p=0.004). However, the amount of sequencing reads required to assess CNA profiles using each approach differed, with CUTseq on average requiring only 10.5% of the sequencing reads needed in the WES analysis (**Figure 3D**).

The regions of the genome enriched for amplifications or deletions across the PreT samples revealed similar patterns between CUTseq and WES CNA calls (**Figure S7**, **Table S10**). Two regions were jointly identified as hotspots for amplifications: 5p15.1 and 8q. The 5p15.1 hotspot encompassed 36 amplified genes including, MYO10 (myosin X) which has been reported to be overexpressed and to drive genomic instability in cancers [77] and contributes to breast cancer aggressiveness and metastasis [78]. Large regions of 8q were enriched for amplifications in both CUTseq and WES analyses (Table S10). The MYC gene is located on this chromosome arm and is commonly amplified in TNBC and promotes immune suppression [79]. Despite the high level of agreement between the two approaches, there were regions of enrichment which were only identified using one approach and not the other. WES identified 3q26.33 as enriched for amplifications, which notably contains PIK3CA which is commonly amplified in TNBC and potentially linked to response to targeted treatment strategies [80]. Alternatively, CUTseq identified a focal amplification of 19q12 harbouring CCNE1 which may confer chemotherapy resistance and has been linked to poor prognosis in TNBC [81], as well as other genes (POP4, PLEKHF1, and TSHZ3) reported to be amplified in cancer [82]. This region of chromosome 19 is adjacent to the centromere which may explain why it was only identified using CUTseq. Similarly, CUTseq identified Xq13.2 as a hotspot for deletions which contains *XIST*, the loss of which has been linked to accelerated tumour growth and brain metastases in breast cancer [83]. Overall, comparisons of WES and CUTseq estimates of CNA, arm level events and ploidy (**Figure 3D**) suggest that these complementary assays provide similar results where they can be compared, while each may still contribute novel information (**Figure S7**). This supports our use of a union of WES/CUTseq CNA calls in the following analyses.

### Pervasive aneuploidies and whole genome duplication drives tumour evolution

Though genome-wide SNV loads show broad stability over time (**Figure 1**), chromosome arm aneuploidies are dynamic, seen abundantly across PreT samples, but often showing dramatic losses in PostT samples (**Figure 4A**, Table S13). These aneuploidies were frequent across all RCB classes and affected almost all chromosomes, with gains of 1q, 8q, 19q and 22q most frequent at PreT (**Figure 4A, 4B**). Dramatic decreases in arm alterations were observed between PreT and PostT samples across most patients, consistent with a disproportionate impact of NACT on tumour lineages carrying aneuploidies. These arm level alterations then often reappeared in later (R) samples but usually involved novel CNA events, suggesting the dominance of distinct lineages in late disease (**Figure 4A**). In general, R samples are notably enriched for these novel events that mimic PreT arm gains. Consistent with timepoint specific advantages to particular tumour clones, certain arm gain events, such as 19q gain seen at PreT, appear to be replaced at later timepoints by 19q arm loss at PostT in multiple patients (**Figure 4A**). This significant pattern of decreased CNA burden at PostT as compared to PreT and R was seen when comparing the mean proportion of the genome altered by amplifications (**Figure 4C**). The prevalence of gain or loss for certain chromosome arms (2p, 7p, 10p, and Xp, Chi-squared p<0.05 with Bonferonni correction) was significantly related to a specific sampling time point (**Figure 4B**). In contrast, certain aneuploidies, particularly gain of 1q and 8q, showed convincing evidence of persistence across the course of treatment, even in PostT samples where most other aneuploidies disappeared (**Figure 4B**). This may suggest ‘aneuploidy addiction’ - where particular aneuploidies are required for tumourigenesis - which has recently been demonstrated *in vitro* in breast cancer cell lines [84]. Strikingly, tumour cell ‘addictions’ to both 1q and 8q gains were found to confer strong advantages in this study, enhancing tumour cell growth, and 1q gains were found to be among the earliest alterations to occur in breast tumour development [84]. However, in this *in vitro* study the mutational mechanisms underlying these aneuploidies remained unknown. The 1q and 8q gains seen here, in TNBC *in vivo*, often occur on a background of pervasive CNA gains seen across the vast majority of chromosomes (**Figure 4B**, **4C**, Table S11), suggesting they may be a result of WGD. Indeed, predicting WGD using a conservative approach (Methods) we found evidence for WGD in 36% (8/22) of patients at PreT, 21% (3/14) at PostT, and in 57% (4/7) at R. All tumours with WGD carry 8q gains, and all but one carry 1q gains at PreT (**Figure 4A**), suggesting that WGD may be exploited to generate advantageous aneuploidies during earlier stages, but also later at recurrence (**Figure 4D**).

**Figure 4.**
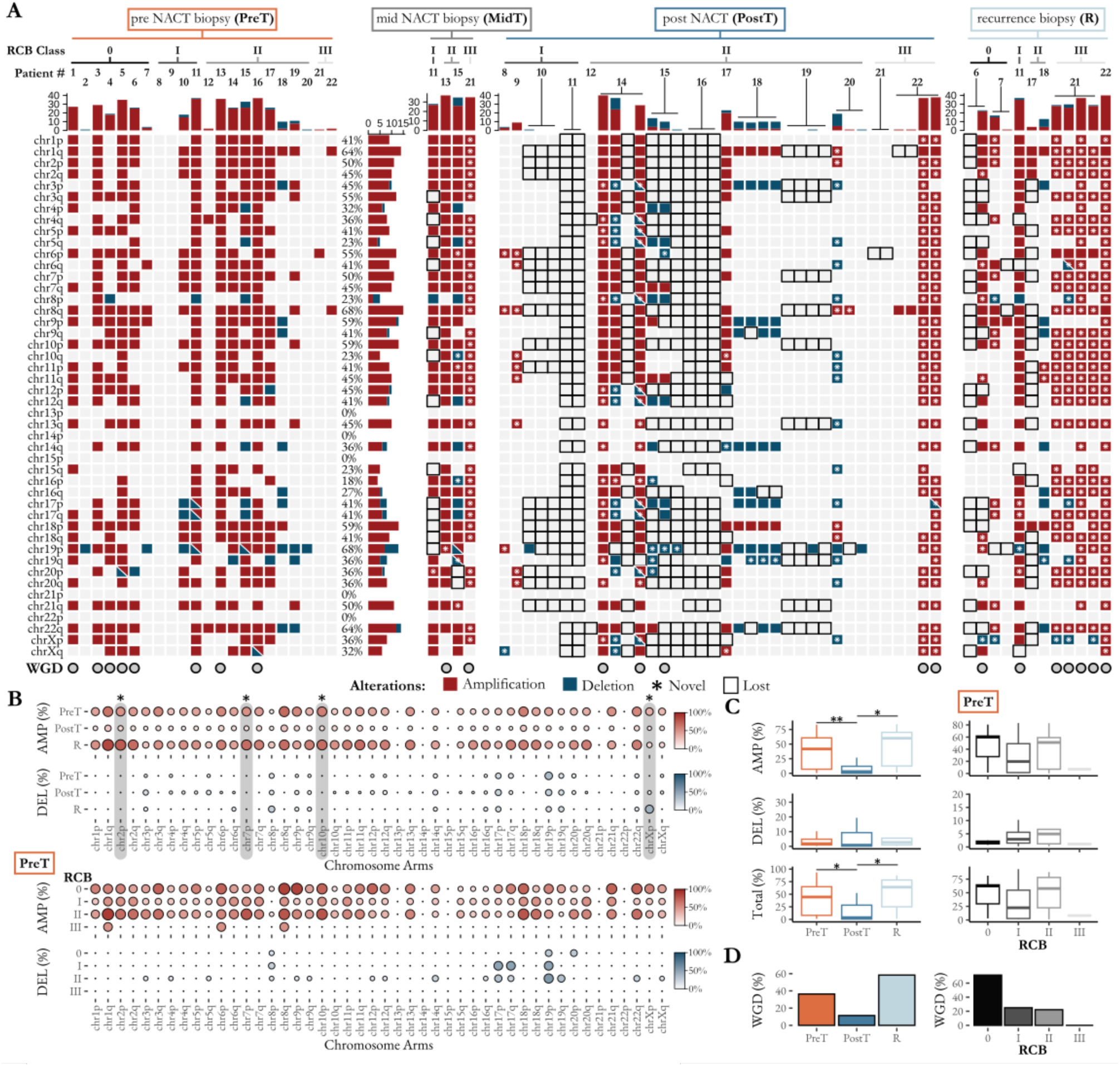
The fate of chromosome arm aneuploidies during NACT. **(A)** The prevalence of aneuploidy for each chromosome arm (rows) longitudionally (four columns representing timepoints: PreT, MidT, PostT and R) are shown for each patient. For PostT and R samples a patient number on top of a T-bar shows the number of multiregional samples assayed from that patient. A chromosome arm was determined to be aneuploid if >60% of the total chromosome arm length was amplified (red) or deleted (blue) by the union CNA calls of WES and CUTseq. Where methods made different calls, the square contains both red and blue. Percentage of the genome with aneuploidies is indicated by bar charts above each column. Aneuploidies lost between PreT and subsequent time points are denoted by a square with a black outline, and newly emerging aneuploidies in MidT, PostT and R, that are not detected in PreT, are indicated with asterisks. Samples estimated to have undergone WGD are indicated with a grey circle. **(B)** The prevalence of chromosome arm aneuploidies at each timepoint (PreT, PostT, R) and for each RCB class (0, I, II III). Size of circles indicate proportion of patients in each category with amplifications (red) and deletions (blue). Grey background and asterisk indicate significant variation in prevalence (amplification or deletion) between time points using Chi-squared tests (p≤0.05 using Bonferonni correction for multiple testing). **(C)** Genomic instability measured as the proportion of each genome amplified (AMP), deleted (DEL) or combination of amplified and deleted (Total) across the time course (left) and RCB class (right). Statistical significance was determined using the Kruskal-Wallis rank sum test, and pairwise Wilcoxon rank sum test. Significance levels are indicated as follows: *p < 0.05; **p < 0.01. **(D)** Proportion of patients with WGD defined as ploidy ≥3. The left panel indicates proportion of patients at each timepoint (PreT, PostT and R) with WGD. The right panel indicates the proportions of patients with WGD at PreT for RCB classes (0, I, II, III).

Gains of 1q and 8q were also seen in 27% (6/22) of PreT samples lacking evidence for WGD, suggesting other mutational mechanisms also produce these alterations. The re-emergence of WGD may reflect the rise in frequency of pre-existing clones with WGD, or further *de novo* WGD events in later disease. This aligns with recent work in ovarian cancer proposing that WGD is an ongoing mutational process promoting evolvability [85]. Importantly, the prevalence of WGD was not uniformly distributed across RCB classes, with the majority (5/7) of patients with the best response to NACT (RCB 0) having tumours with WGD at PreT (**Figure 4D**). One of the drugs included in NACT (docetaxel) has been shown to induce polyploidization and aberrant aneuploidies by stabilizing microtubules [86]. The increased chromosome instability introduced by NACT may cause tumour cells that have previously undergone WGD to become unviable leading to improved treatment response and reduced residual cancer burden.

Given the large changes in ploidy and frequent WGD seen during TNBC evolution, many genes undergo recurrent alterations (**Figure 5**). Among the genes amplified are many that have previously been reported to undergo alterations during progression (**Figure 5A**) in prior studies of the TNBC genome [8,49,62]. Among the most recurrently amplified are *MYC* (located at 8q24.21), *AKT3* (1q43) and *MDM4* (1q32.1). All three amplifications are present in the majority of PreT samples, are very rarely deleted, and also show evidence of persistence in multiple patients throughout treatment and into recurrence (**Figure 5A, B**), consistent with the 1q and 8q chromosome arm gains that also persist through treatment (**Figure 4A**). All three have been specifically linked to TNBC via their upregulation and amplification. *MYC* amplification is commonly observed in TNBC and is thought to suppress innate immunity and allow tumour immune escape [79]. *AKT3* amplification has been observed in multiple breast cancer subtypes, including TNBC, and is thought to mediate resistance to tamoxifen [87]. *MDM4* has emerged as an important factor in promoting TNBC tumour formation and metastasis, via its role as a *TP53* inhibitor [62]. It therefore seems plausible that the frequent and persistent 1q and 8q arm gains seen in these tumours (**Figure 4A**) result in increases in the expression of one or more of these genes, though we lack direct evidence for this. This would also offer a potential explanation for tumour cell ‘addiction’ to 1q and 8q gains seen in recent *in vitro studies* [84]. Analysis of altered oncogenic pathways revealed an intricate pattern of synergy between genes in the RTK/ RAS pathway and 1q and 8q gains. Amplifications of eight genes (**Figure 5A, B**) in the RTK/ RAS pathway (*EGFR*, *IGF1R*, *KRAS*, *KIT*, *NF1*, *FGFR1*, *PDGFRA* and *MAP3K1*) showed co-occurrence with gains in 1q (12/16 patients) or 8q (14/15).

**Figure 5.**
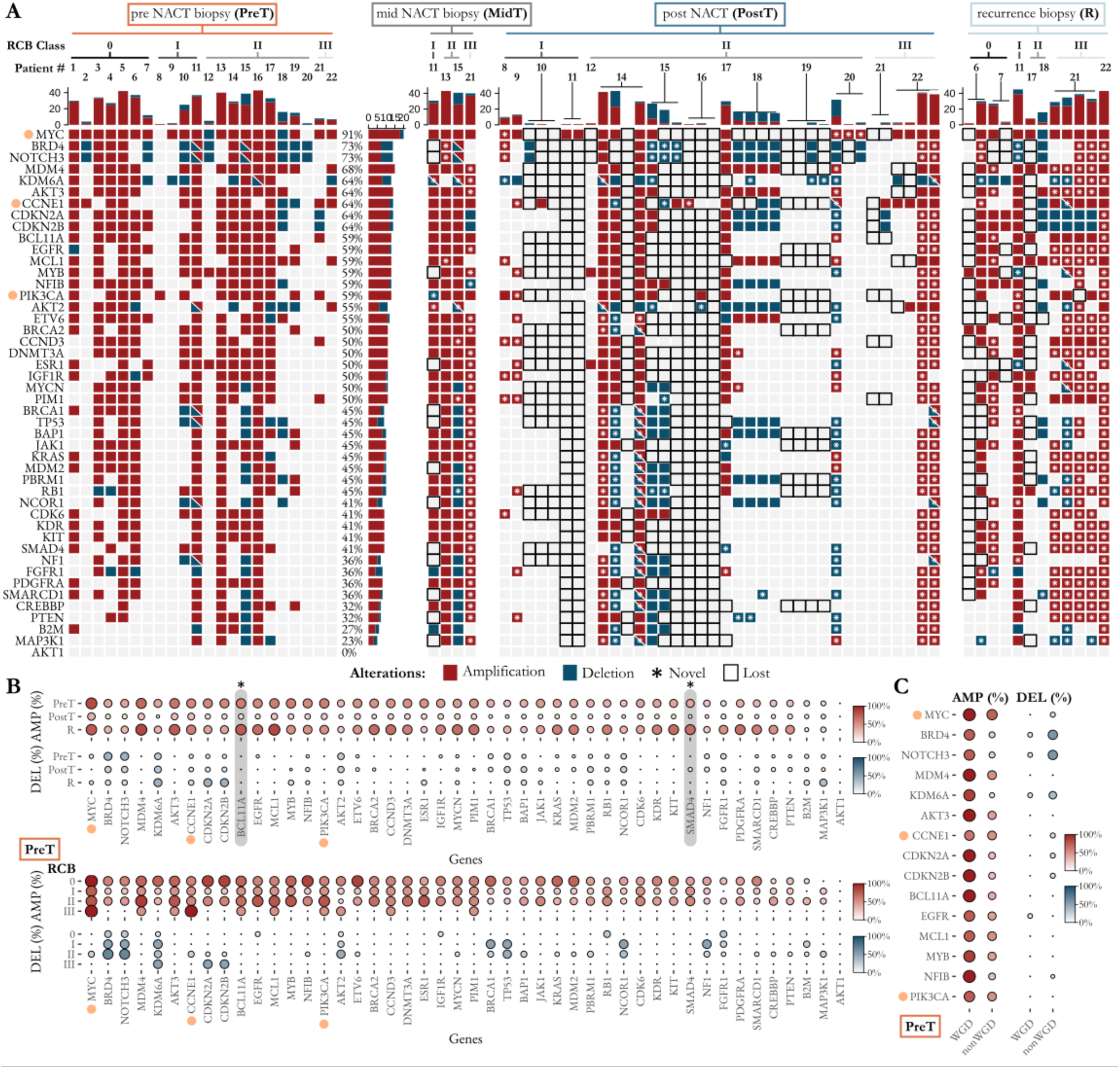
Genes with recurrent CNAs during NACT. **(A)** Oncoplot indicating prevalence of amplification (red) and deletion (blue) of genes previously associated with TNBC progression. Alterations lost between PreT and subsequent time points are denoted by a square with a black outline and newly emerging alterations, in MidT, PostT and R, that are not detected in PreT, are indicated with asterisks. Squares showing both red and blue indicate where WES and CUTSeq disagree over the presence of a newly emerging CNA. RCB class and TNBC patient number are shown along the top, a patient number on top of a T-bar shows the number of multiregional samples assayed from that patient. Orange dots in **(A)**, **(B)** and **(C)** indicate genes identified in CNA hotspots in enrichment analysis (Table S10). **(B)** Prevalence of gene alterations at each timepoint (PreT, PostT, R) and across RCB classes (0, I, II III). Size of circles indicate proportion of patients in each category with gene amplifications (red) and deletions (blue) and a grey background and asterisk indicate significant variation in prevalence (gene amplification or deletion) between time points using Chi-squared tests (p≤0.05 using Bonferonni correction for multiple testing) **(C)** Prevalence of amplification (red) and deletion (blue) in the top 15 altered genes identified in panel A, split according to WGD status of PreT samples.

Other frequently altered genes suggest a more complex mutational landscape. For example, *BRD4* (19p13.12), *NOTCH3* (19p13.2) and *CCNE1* (19q12) undergo amplification and deletion in different samples and at different timepoints (**Figure 5A, B**). This volatile state may suggest the presence of complex rearrangements such as chromothripsis, which has been reported to be particularly enriched on chromosome 19 in high grade serous ovarian cancer [88]. We also observe frequent amplifications and deletions of *KDM6A*, with amplifications common in PreT but deletions at later stages (**Figure 5A, B**), consistent with reports that *KDM6A* loss can reduce sensitivity to the therapeutic agents used to treat TNBC patients [89].

A closer examination of the relationship between gene copy number alterations and WGD status in PreT samples revealed distinct patterns (**Figure 5C**). In samples that had undergone WGD, amplifications of the top 15 most commonly altered genes were highly prevalent (87.5%). In contrast, non-WGD samples showed a lower frequency of amplification (38.6%). Deletions were relatively rare in WGD samples; however, in non-WGD samples, deletions affecting BRD4 and NOTCH3 at 19p13.12 were common, occurring in 57.1% of cases compared to just 12.5% in WGD samples. These findings suggest that WGD shapes genome-wide mutational patterns and drives distinct combinations of candidate driver gene alterations.

### Clinical correlates of mutational loads and aneuploidy

A number of clinically relevant features were found to be inter-correlated (**Figure 6A**). TMB in PreT samples positively correlated with metrics of genomic instability across various scales: CUTseq derived ploidy (R=0.59, p=0.04), total structural variant burden (R=0.73, p=0.0001) and microsatellite instability (R=0.73, p=0.009). As anticipated, the amount of the genome amplified by CNAs correlated with CUTseq (R=0.71, p=0.004) and WES derived ploidy estimates (**Figure 6A**, R=0.73, p=0.0009), and TIL infiltration and RCB score were negatively correlated (R=-0.56, p=0.045) (**Figure 6A**, Figure S1). Investigating relationships between mutations, clinical features and overall patient overall survival (OS) showed that patient OS varied significantly, with patients in RCB class II/III having poorer OS than RCB 0/I patients (p=0.039) (**Figure 6B**). Although HRD scores did not differ significantly between RCB classes (Figure S8), as expected, patients with tumours showing evidence for HRD in PreT samples (defined as scarHRD scores >70) had significantly better OS (**Figure 6B**, p=0.01). This is in agreement with previous studies demonstrating higher HRD scores are associated with a better OS and better sensitivity to treatments such as PARP inhibitors and platinum-based chemotherapy [90,91]. Using a Bayesian approach (Methods) we looked for pairwise co-occurrence or mutual exclusivity among mutational events, including the presence or absence of SNVs in driver genes (**Figure 2**), chromosome arm aneuploidies (**Figure 4**) or CNAs recurrently impacting genes (**Figure 5**) (see Figure S9). The output was dominated by pairwise co-occurrence events between amplifications in genes and chromosome arms, as expected with frequent WGD events (Figure S10).

**Figure 6.**
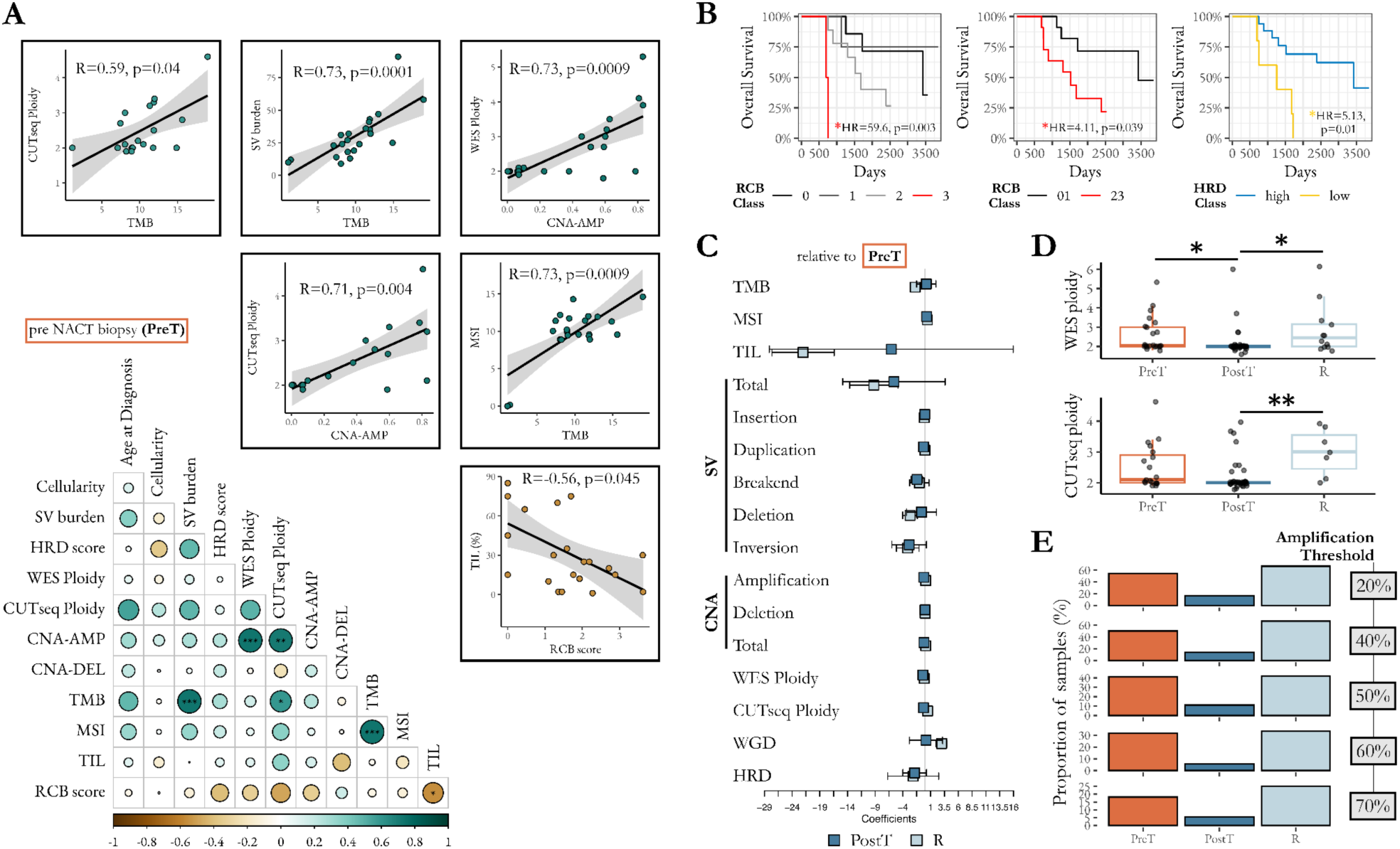
Correlations among mutational and clinical variables. **(A)** Pairwise correlations between genomic alterations and other features of interest in PreT samples, indicated by an all-against-all correlation matrix with strength of correlation indicated by size of coloured circles (bottom left). Additional scatter plots depict the relationships between selected variables showing significant correlations. **(B)** Kaplan-Meier curves depicting relationship between RCB class, HRD score and patient OS. Hazard ratios (HR) and p-values are derived from Cox Proportional-Hazard models. **(C)** Forest plot of coefficients derived from LMM (for continuous variables) or GLMM (for binary variables) for the complete trajectory of each molecular or clinical feature over time, indicating features increasing (positive coefficients) or decreasing (negative) over time. Coefficients for PostT (blue) and recurrence (light blue) are shown relative to PreT samples. **(D)** Ploidy estimates from ASCAT.sc for each sample across time using WES and CUTseq. Median ploidy estimates for WES and CUTseq respectively, were PreT: 2.05 and 2.1, PostT: 2.0 and 2.0 and R: 2.45 and 3.0. Asterisks indicate statistical significance using Pairwise Wilcoxon Rank Sum Tests (adjusted p-values <0.01=**, <0.05=*). **(E)** The proportions of samples (PreT, PostT or R) showing evidence of WGD, based upon various thresholds for the mean fraction of genome amplified across WES and CUTseq approaches.

To analyse longitudinal patterns within each individual patient, we employed Linear Mixed Model (LMM) and Generalised LMM (for binary dependent variables) modelling approaches for all features. The modelling indicates that several molecular and clinical features change significantly across time points (**Figure 6C**, Table S14), showing patterns that are broadly consistent with the analyses based upon pairwise comparisons between timepoints (**Figure 4**). For instance, from PreT to R, we observe a decrease in TILs (*β* = −22.03, SE=5.62, p<0.0001) and a reduction of SV burden (*β* = −9.24, SE=4.38, p=0.035). Similarly, the odds of WGD shrinks to 18% of the initial odds from PreT to PostT (p=0.038), indicating that WGD is significantly less likely to occur at the PostT time point compared to PreT (Table S14). WGD is also commonly seen in R samples (**Figure 4D**), which is likely to reflect the resurgence of WGD bearing clones in these recurrent tumours. In fact, even though median ploidy is greater than 2 in PreT samples, median (WES or CUTseq) ploidy is higher at recurrence, suggesting multiple WGD events in some samples (**Figure 6D**). As the threshold for WGD (mean fraction of genome amplified over WES and CUTseq datasets) is increased from 20% to 70%, PreT and R samples were consistently higher than PostT samples, demonstrating that this trend is not dependent on the threshold by which WGD was determined (**Figure 6E**).

We conclude that WGD events are an under-appreciated mutational mechanism underlying the evolution of TNBC during treatment. These events appear to drive the acquisition of advantageous aneuploidies, particularly 1q and 8q gains, and are predicted to impact known cancer driver genes. WGD bearing tumours are common at PreT and presumably confer proliferative advantages before treatment, but these tumours also appear to be unusually sensitive to NACT, resulting in a strong decline in the ploidy of PostT samples. At later stages higher ploidy mediated by WGD, again appears to offer advantages to dominant clones within R samples.

## Discussion

Using two complementary sequencing approaches (WES and CUTseq), we have reconstructed the mutational landscape of TNBC and tracked its dynamics during NACT. The genomic and transcriptomic landscape of TNBC has been examined in several previous studies [6–8,51], but TNBC temporal mutational dynamics during chemotherapy treatment is poorly studied. Here, we describe a cohort with longitudinal sampling, providing an unusual opportunity to determine how the mutational landscape responds to treatment, including SNVs, CNAs and aneuploidies. We found that although the genome-wide SNV mutational burden remained stable during disease progression, the loss and emergence of SNVs in known TNBC driver genes was common in response to treatment. These complex dynamics were seen in *TP53*, *MICA*, *CYP2D6*, *BRCA1* and *BRCA2*, suggesting clonal loss and replacement events over time, consistent with selection favouring different variants at different times in the same driver gene.

Dramatic genome-wide structural shifts were seen in all samples, reflected in CNA and chromosome arm aneuploidies, with a substantial bias to duplications. Overall, the abundant structural variation seen in PreT samples was followed by frequent loss of these alterations after treatment, but the re-emergence of similar alterations at recurrence. These alterations include advantageous aneuploidies increasing tumour growth *in vitro*, such as 1q gain, that has been proposed to drive tumourigenesis via an increase in *MDM4* expression, which phenocopies the effects of *TP53* inactivating mutations to suppress *TP53* signalling [84]. Such aneuploidies have also been demonstrated to create collateral therapeutic vulnerabilities via the toxic effects of duplicating many 1q genes, which can represent novel drug targets [84]. We also see positive correlations between ploidy and TMB, suggesting tumours undergoing WGD may present higher neoantigen loads, targetable by neoantigen vaccines, underlining the promise of such emerging approaches in TNBC [92].

Relatively early, and frequent (36% of PreT samples) WGD events appear to drive or at least co-occur with the extensive aneuploidies seen pre-treatment, consistent with a recent study of TNBC patient-derived xenograft samples that lacked longitudinal sampling [20]. Notably a higher frequency of PreT WGD was seen in patients with the best response to NACT (RCB0/1: 55%; RCB2/3: 18%), suggesting that early WGD may confer susceptibility to treatment. WGD was also frequently found at recurrence (58% of R samples), either reflecting the re-emergence of proliferative WGD bearing clones, or late, *de novo* WGD events. To our knowledge this is the first evidence for WGD as a common event in both pre-treatment and recurrent TNBC tumours. Subsequent studies with larger sample sizes and generating WGS data will be better placed to make more detailed estimates of the impact of WGD on clinical outcomes.

These unusual, longitudinally sampled data suggest the presence of WGD itself may be a useful prognostic biomarker of NACT response, associated with better or worse outcomes, depending on the time of sampling. Emerging studies also suggest that WGD may be a targetable vulnerability [93,94]. Detecting WGD and genome-wide CNA accurately in a FFPE tumour sample is not possible using the most widely used profiling technologies (targeted sequencing panels) while WES and WGS are not financially viable options for many healthcare providers. However, we have demonstrated that CUTseq represents an affordable assay that provides accurate results in the setting of structurally complex tumour genomes sampled using FFPE blocks and primary tumour sections.

## Conclusions

Determining the origins and dynamics of structural complexity in tumourigenesis over time is critical to understand the genomic heterogeneity between patients and within tumours. In this study we reveal the genome-wide shifts in structural variation that underlie TNBC evolution during treatment. We present evidence that WGD in pre-treatment biopsies is associated with better response to NACT, but also appears to confer advantages to tumours at recurrence. Validation in larger, temporally sampled cohorts is required to confirm these results, but the analysis of large numbers of samples will rely on the development of affordable sequencing strategies that can be used to profile the (FFPE) samples that are generally available from healthcare providers. We have demonstrated that CUTseq and WES can be combined to that end, and readily provide new insights into tumour evolution over time.

## Supplementary information

Additional file 1: a PDF file containing all supplementary figures (Fig. S1–S10).

Additional file 2: an Excel table containing all supplementary tables (Table S1–S14).

## Acknowledgements

The authors thank the patients and their families for their participation in this academic study. We are also grateful for the time and efforts of the breast radiologists and surgeons and nurses of the Edinburgh Breast Unit for collecting samples. The authors would like to acknowledge the Edinburgh Clinical Research Facility for the DNA sequencing and support, and the NRS Lothian Bioresource for additional tissue samples. Some of the results published here are in part based upon data generated by the TCGA Research Network: https://www.cancer.gov/tcga.

## Authors’ contributions

O.O., F.S., A.E. and C.A.S. designed the study and supervised analysis of the cohort. F.S. performed laboratory work and D.P.B. performed the bioinformatic analyses. A.M., M.I.O. and A.E. provided additional analysis. A.I. performed histopathological review. O.O designed the NEO and metcfDNA studies, arranged relevant ethical approvals and recruited patients (under her care throughout their breast cancer journey). N.W collected tissue samples and gathered patient clinicopathological characteristics, from national health electronic patient records. D.P.B., F.S., O.O. and C.A.S. wrote the manuscript, which was read and approved by all authors.

## Data availability

Genomic data generated in this study have been submitted to the European Genome/ Phenome Archive, with accession numbers EGAD00001015684 (CUTseq data) and EGAD00001015687 (WES data).

## Declarations

### Ethics approval and consent to participate

Collection of human tissues and relevant clinicopathological data were approved by the South East Scotland Research Ethics Committee 01 with REC references: 13/SS/0236 and 15/ES/0094, and all patients consented to sample collection and use of their data in research

### Consent for publication

Not applicable

