## Supplementary Figures S1-S10 for "Frequent Whole-Genome Duplication Events Drive the Genomic Evolution of Triple-Negative Breast Cancer During Neoadjuvant Chemotherapy"

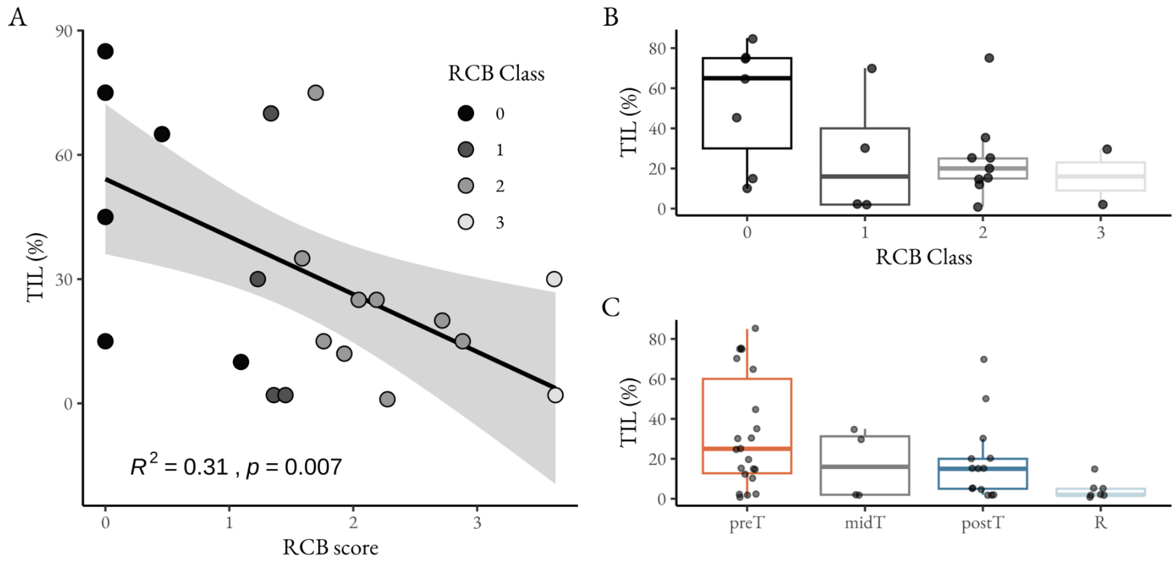

**Figure S1 | Tumour-infiltrating lymphocytes (TILs) in TNBC during treatment. (A)** The linear relationship between TILs and RCB. The nodes are coloured according to the patient's determined RCB class. **(B)** The distribution of TILs sorted by RCB Classes. **(C)** The distribution of TILs sorted by the sampling timepoints.

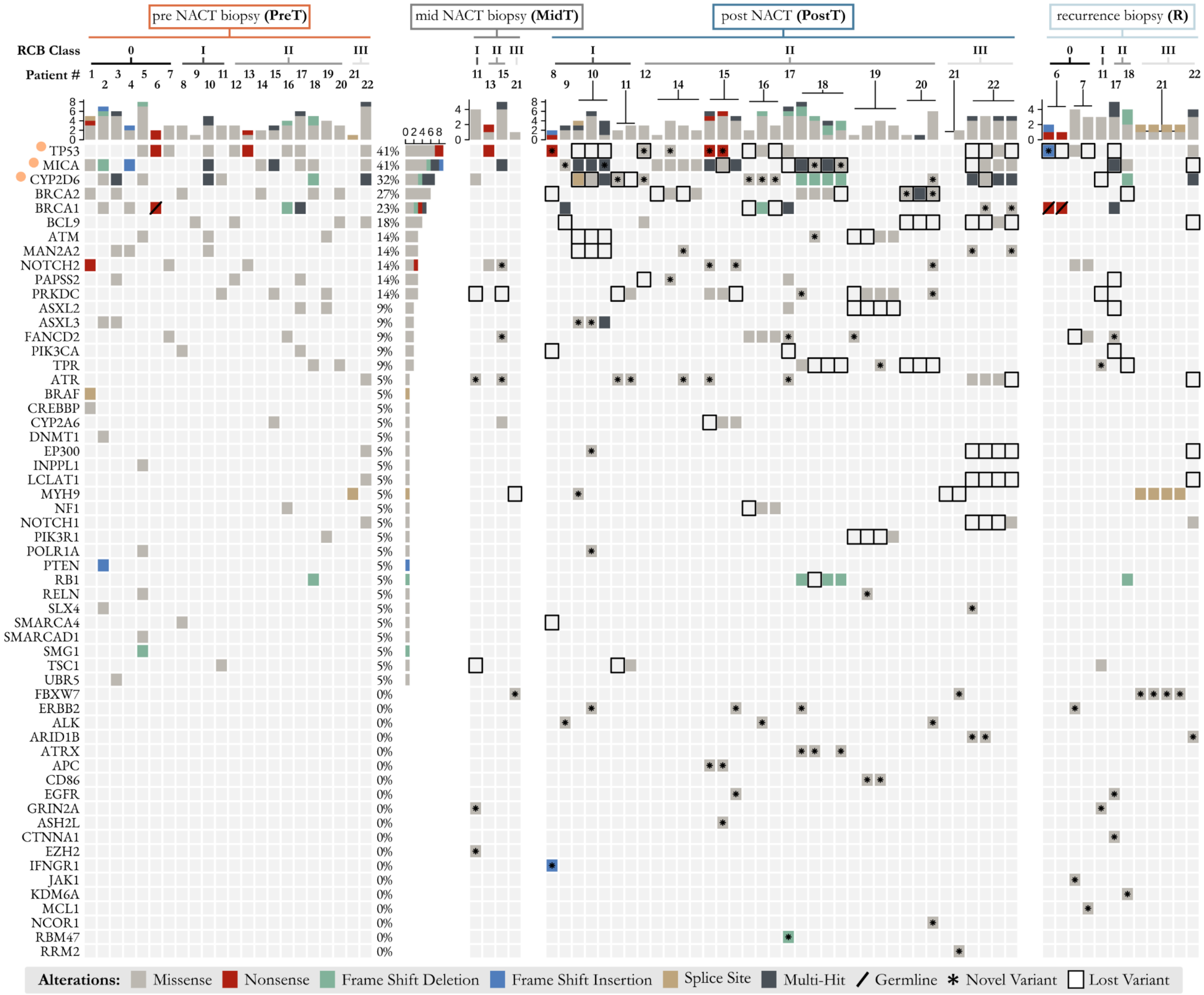


**Figure S2 | The complete SNV landscape of TNBC genes within our patient cohort.** Oncoplot indicating the presence or absence of mutational variants in coding regions of genes. Variants detected for each TNBC patient (patient number indicated at the top) are shown at each timepoint, with variant type indicated by a coloured square (see key, bottom). For PostT and R samples a patient number on top of a T-bar shows the number of multiregional samples assayed from that patient. Variants lost between PreT and PostT or between PreT and R, are denoted by a square with a black outline. Variants not found in PreT that are detected in PostT or R, are denoted by an asterisk. Orange dots indicate genes identified as driver candidates and germline variants are indicated with a solid diagonal line. Where multi-region samples exist for the same timepoint they generally support the existence of the same driver mutation.

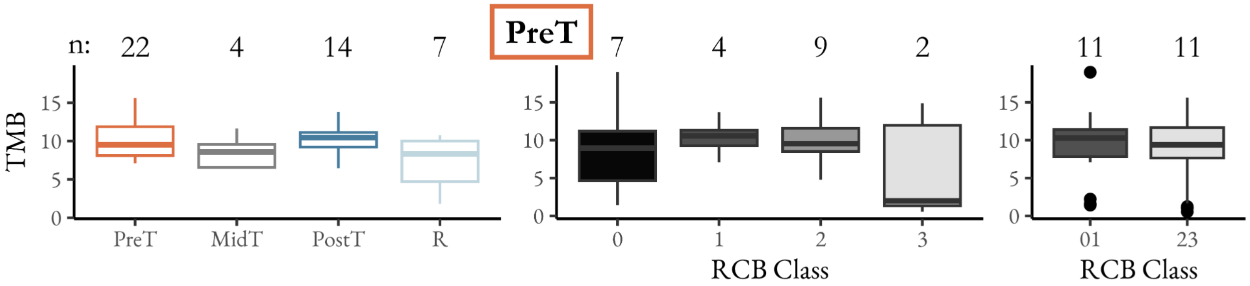


**Figure S3 | Tumour mutational burden (TMB) distributions in PreT samples.** TMB calculated as SNVs per Mb for each patient across timepoints, orange = PreT, grey = MidT, blue = PostT, light blue = R; and RCB classes 0, I, II and III and combined 0/I compared to II/III. The number of patients in each group is indicated along the top.

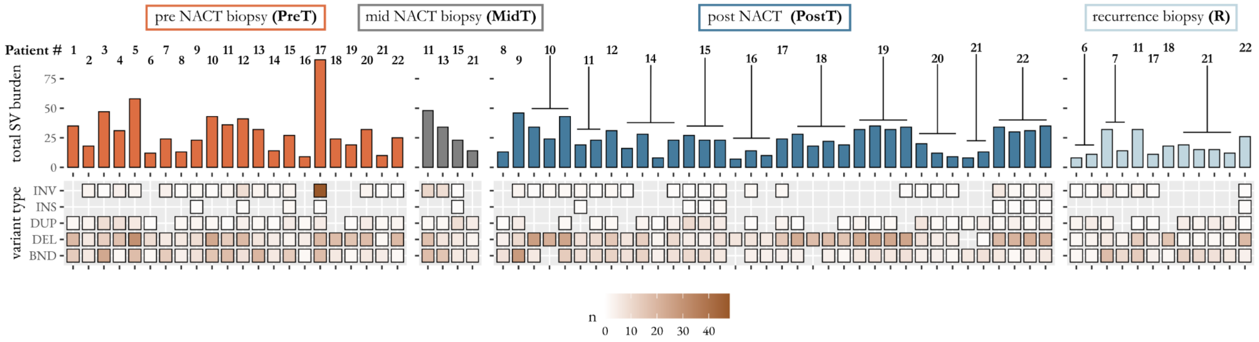


**Figure S4 | The longitudinal structural variant (SV) landscape of TNBC.** The total number of SVs detected in each sample are shown in the barplot, patient numbers shown at the top, with T bars indicating samples where multiregional samples were assayed from that patient. The heatmap indicates the contribution of each variant type (inversion, insertion, duplication, deletion, breakend) for each patient at each timepoint.

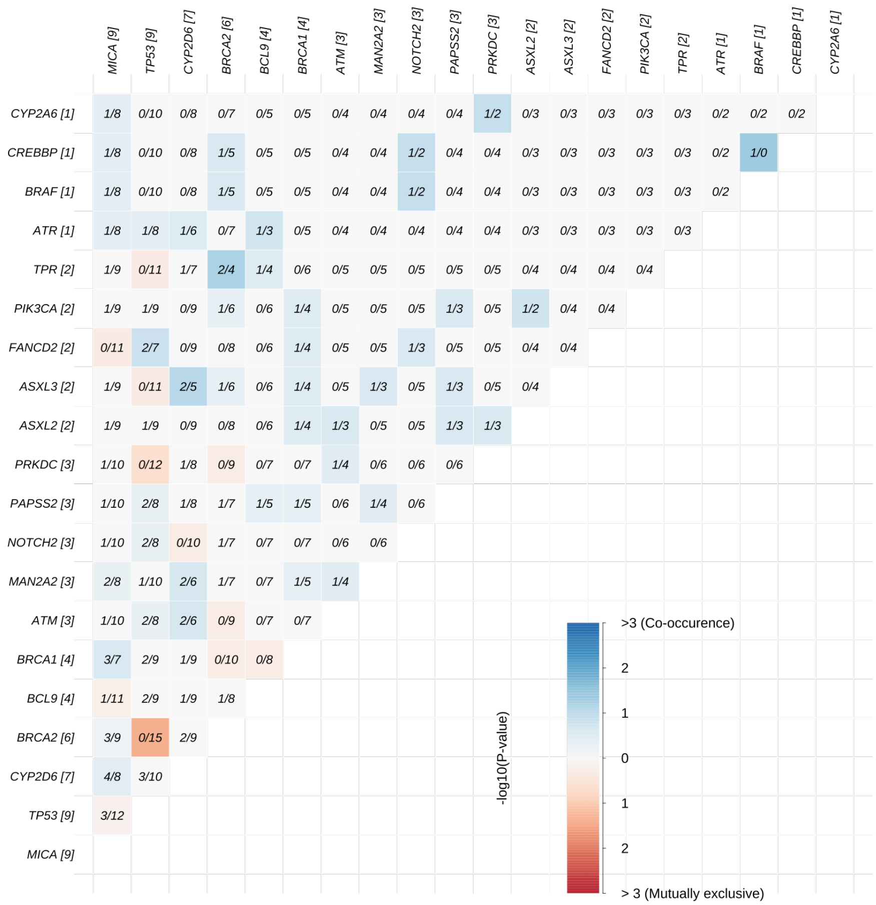


**Figure S5 | Presence/ Absence matrix of somatic SNVs.** Pairwise mutual exclusivity (red) or co-occurrence (blue) among somatic mutations in PreT samples.

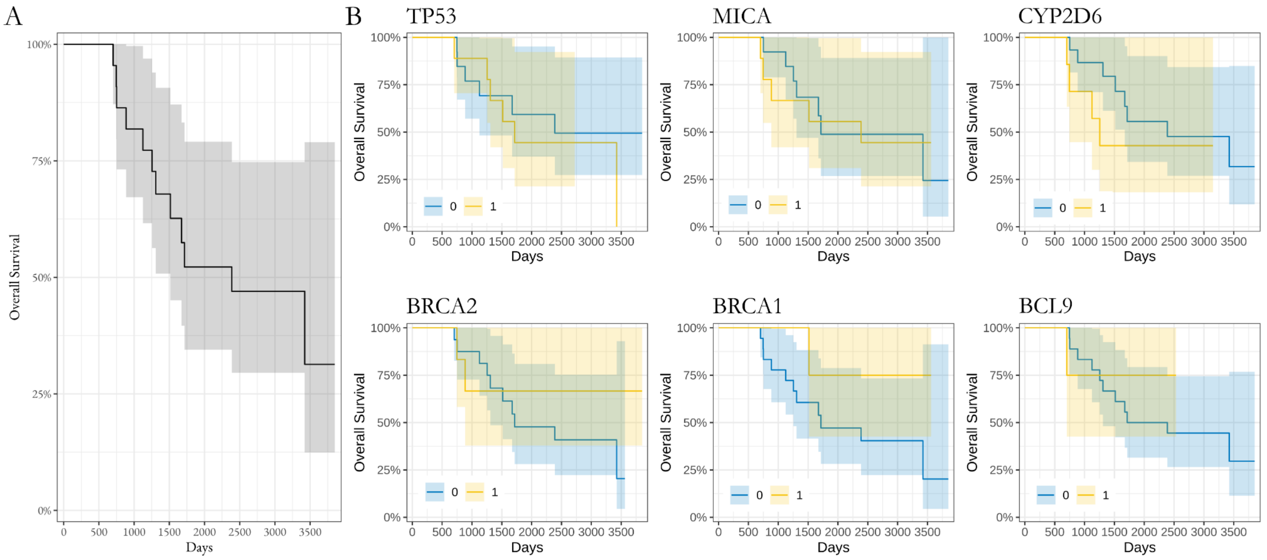


**Figure S6 | Mutational effects on patient survival. (A)** Kaplan-Meier survival curve of patient overall survival. **(B)** Kaplan-Meier survival curve based on the presence (yellow) or absence (blue) of SNV variants in the top 6 genes (TP53, MICA, CYP2D6, BRCA2, BRCA1 and BCL9).

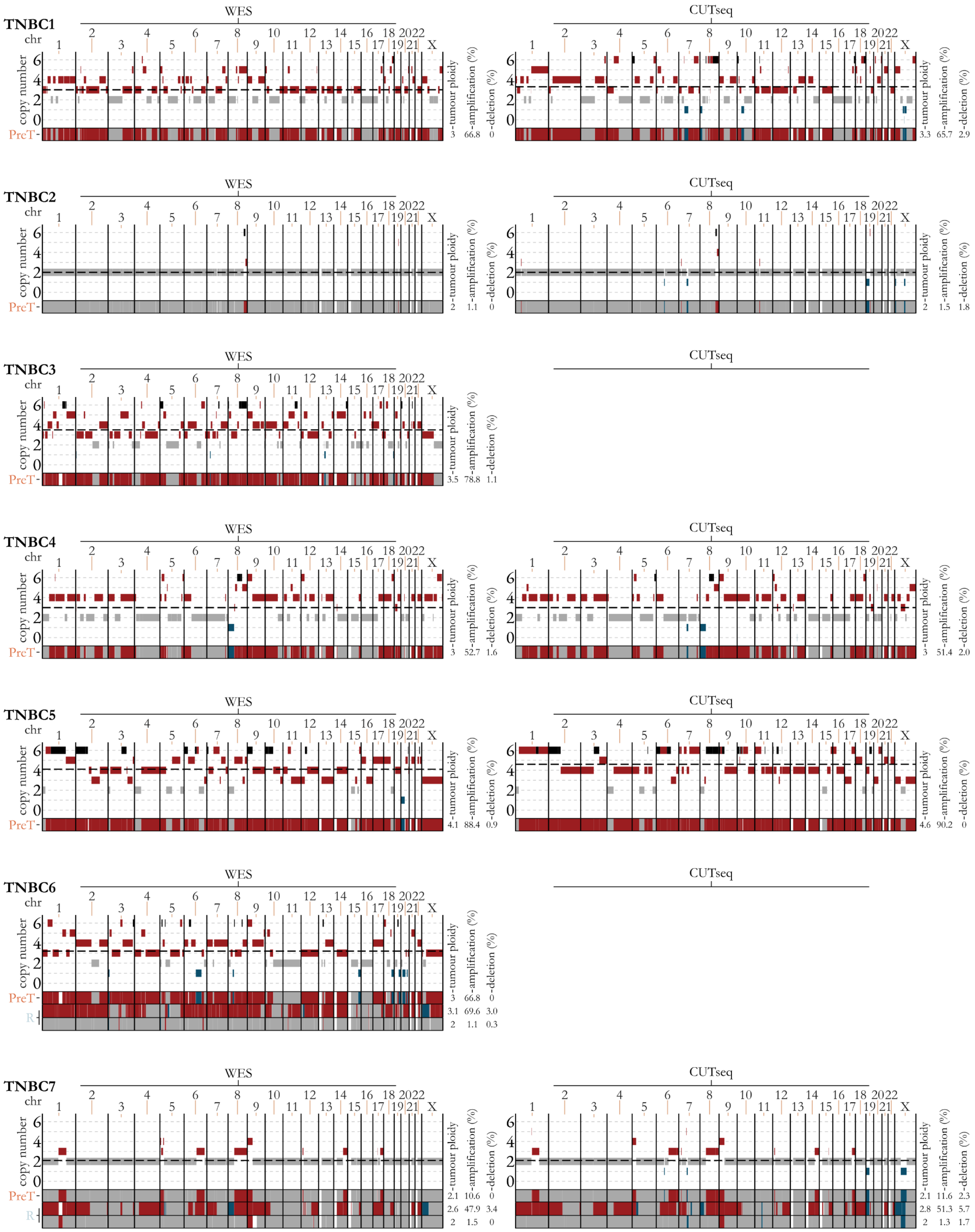

TNBC1
WES
CUTseq
chr
copy number
PreT
tumour ploidy
amplification (%)
deletion (%)
3 66.8 0
3.3 65.7 2.9
TNBC2
2 1.1 0
2 1.5 1.8
TNBC3
3.5 78.8 1.1
TNBC4
3 52.7 1.6
3 51.4 2.0
TNBC5
4.1 88.4 0.9
4.6 90.2 0
TNBC6
R
3 66.8 0
3.1 69.6 3.0
2 1.1 0.3
TNBC7
2.1 10.6 0
2.6 47.9 3.4
2 1.5 0
2.1 11.6 2.3
2.8 51.3 5.7
2 1.3 1.7

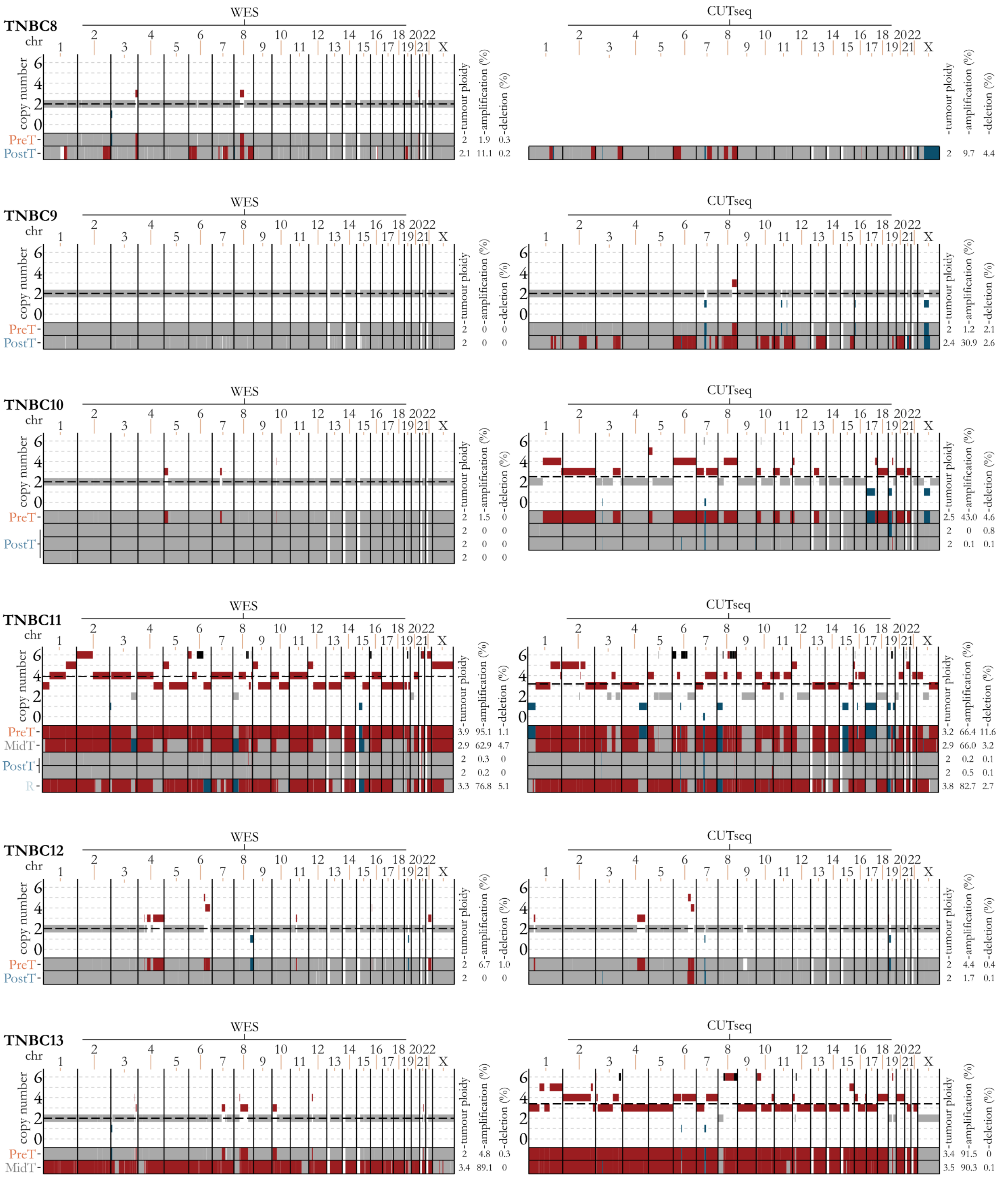
TNBC8
WES
CUTseq
chr
copy number
PreT
PostT
tumour ploidy
amplification (%)
deletion (%)
WES PreT 2 1.9 0.3
WES PostT 2.1 11.1 0.2
CUTseq PostT 2 9.7 4.4
TNBC9
WES PreT 2 0 0
WES PostT 2 0 0
CUTseq PreT 2 1.2 2.1
CUTseq PostT 2.4 30.9 2.6
TNBC10
WES PreT 2 1.5 0
WES PostT 2 0 0
WES PostT 2 0 0
WES PostT 2 0 0
CUTseq 2.5 43.0 4.6
CUTseq 2 0 0.8
CUTseq 2 0.1 0.1
TNBC11
MidT
R
WES PreT 3.9 95.1 1.1
WES MidT 2.9 62.9 4.7
WES PostT 2 0.3 0
WES PostT 2 0.2 0
WES R 3.3 76.8 5.1
CUTseq 3.2 66.4 11.6
CUTseq 2.9 66.0 3.2
CUTseq 2 0.2 0.1
CUTseq 2 0.5 0.1
CUTseq 3.8 82.7 2.7
TNBC12
WES PreT 2 6.7 1.0
WES PostT 2 0 0
CUTseq 2 4.4 0.4
CUTseq 2 1.7 0.1
TNBC13
WES PreT 2 4.8 0.3
WES MidT 3.4 89.1 0
CUTseq 3.4 91.5 0
CUTseq 3.5 90.3 0.1

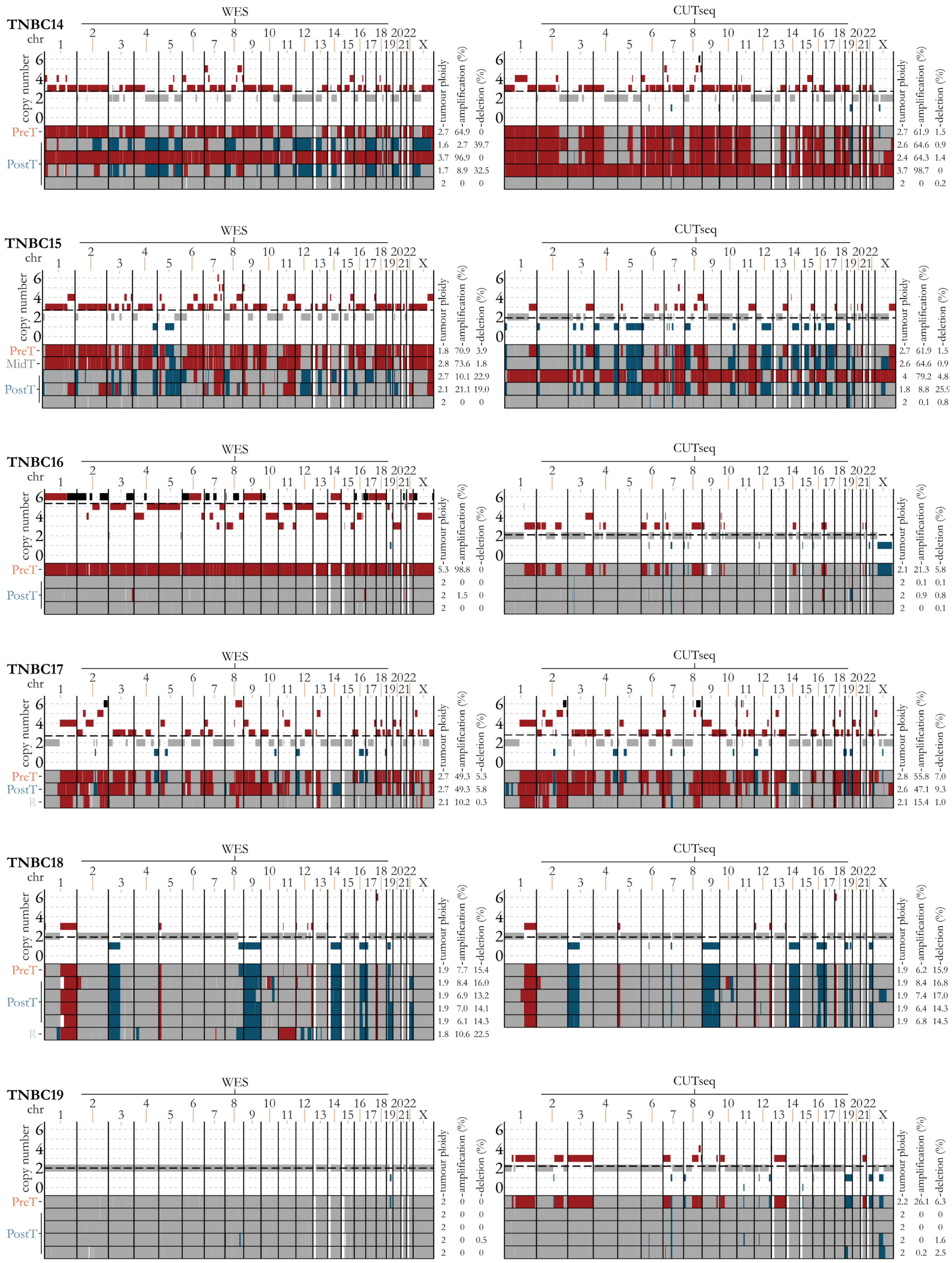
TNBC14
WES
CUTseq
chr
copy number
PreT
PostT
tumour ploidy
amplification (%)
deletion (%)
2.7 64.9 0
1.6 2.7 39.7
3.7 96.9 0
1.7 8.9 32.5
2 0 0
2.7 61.9 1.5
2.6 64.6 0.9
2.4 64.3 1.4
3.7 98.7 0
2 0 0.2
TNBC15
PreT
MidT
PostT
1.8 70.9 3.9
2.8 73.6 1.8
2.7 10.1 22.9
2.1 21.1 19.0
2 0 0
2.7 61.9 1.5
2.6 64.6 0.9
4 79.2 4.8
1.8 8.8 25.9
2 0.1 0.8
TNBC16
5.3 98.8 0
2 0 0
2 1.5 0
2 0 0
2.1 21.3 5.8
2 0.1 0.1
2 0.9 0.8
2 0 0.1
TNBC17
PreT
PostT
R
2.7 49.3 5.3
2.7 49.3 5.8
2.1 10.2 0.3
2.8 55.8 7.0
2.6 47.1 9.3
2.1 15.4 1.0
TNBC18
1.9 7.7 15.4
1.9 8.4 16.0
1.9 6.9 13.2
1.9 7.0 14.1
1.9 6.1 14.3
1.8 10.6 22.5
1.9 6.2 15.9
1.9 8.4 16.8
1.9 7.4 17.0
1.9 6.4 14.3
1.9 6.8 14.5
TNBC19
2 0 0
2 0 0
2 0 0
2 0 0.5
2 0 0
2.2 26.1 6.3
2 0 0
2 0 0
2 0 1.6
2 0.2 2.5

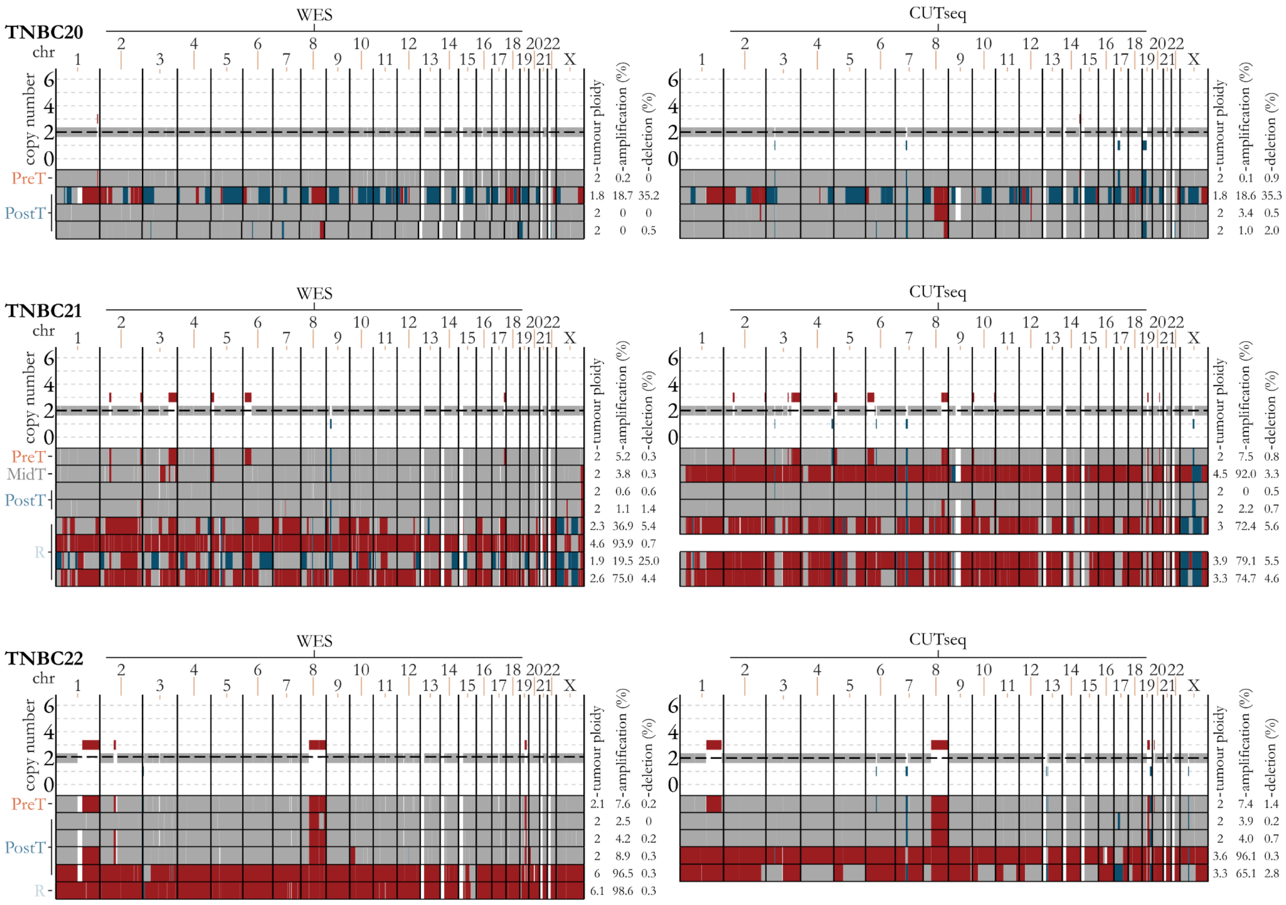


**Figure S7 Longitudinal CNA calls for each patient indicating the consistent patterns exhibited between WES and CUTseq.** WES plots for each patient are on the left, CUTseq (where performed) on the right, with TNBC patient numbers indicated at the top left of each WES plot. Amplified genomic regions are shown in red, deleted regions in blue, diploid regions in grey and white indicates regions where copy number was unobtainable. The dashed black line indicates the estimated tumour ploidy. Each individual plot shows the estimated copy number (y-axis) for each chromosome (x-axis) for the PreT sample. Below this, amplifications and deletions are summarised for all timepoints, including multiregional samples, as indicated on the left. Estimated tumour ploidy, and the percentage of the genome either amplified or deleted from each approach is shown to the right of each plot.

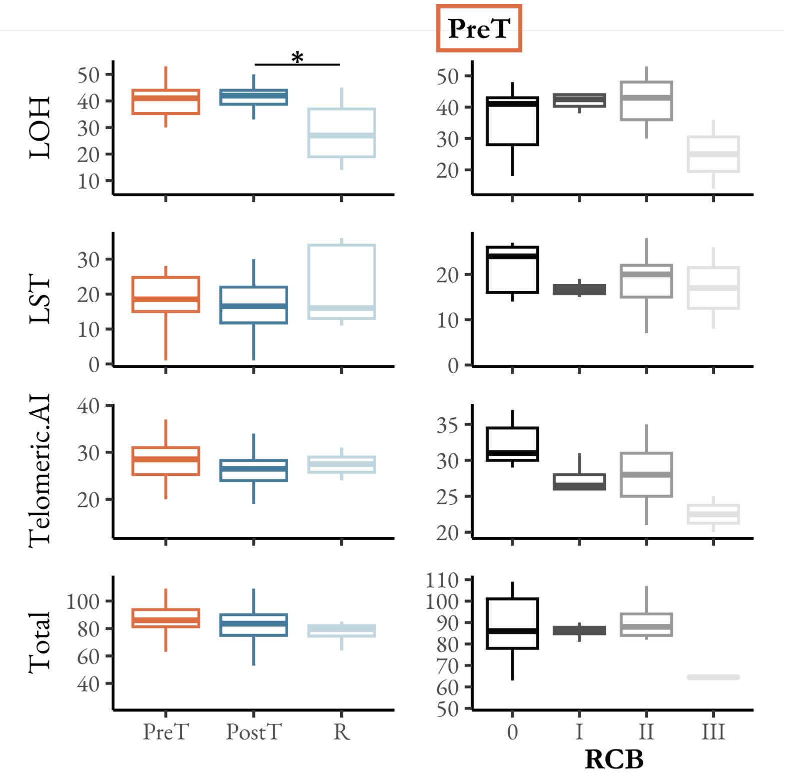


**Figure S8 | Proportion of samples showing evidence for HRD over time and RCB classes.** HRD was estimated using scarHRD [64] which integrates three scores from patterns of loss of heterozygosity (LOH), large scale transitions (LST) and telomeric allelic imbalances (Telomeric.AI). The LOH score is defined as the number of LOH regions exceeding 15 Mb, LST score as the number of large-scale transitions exceeding 10 Mb, and Telomeric.AI score as the number of telomeric allelic imbalances. Total represents the overall likelihood of HRD based upon the combination of LOH, LST and Telomeric.AI scores.

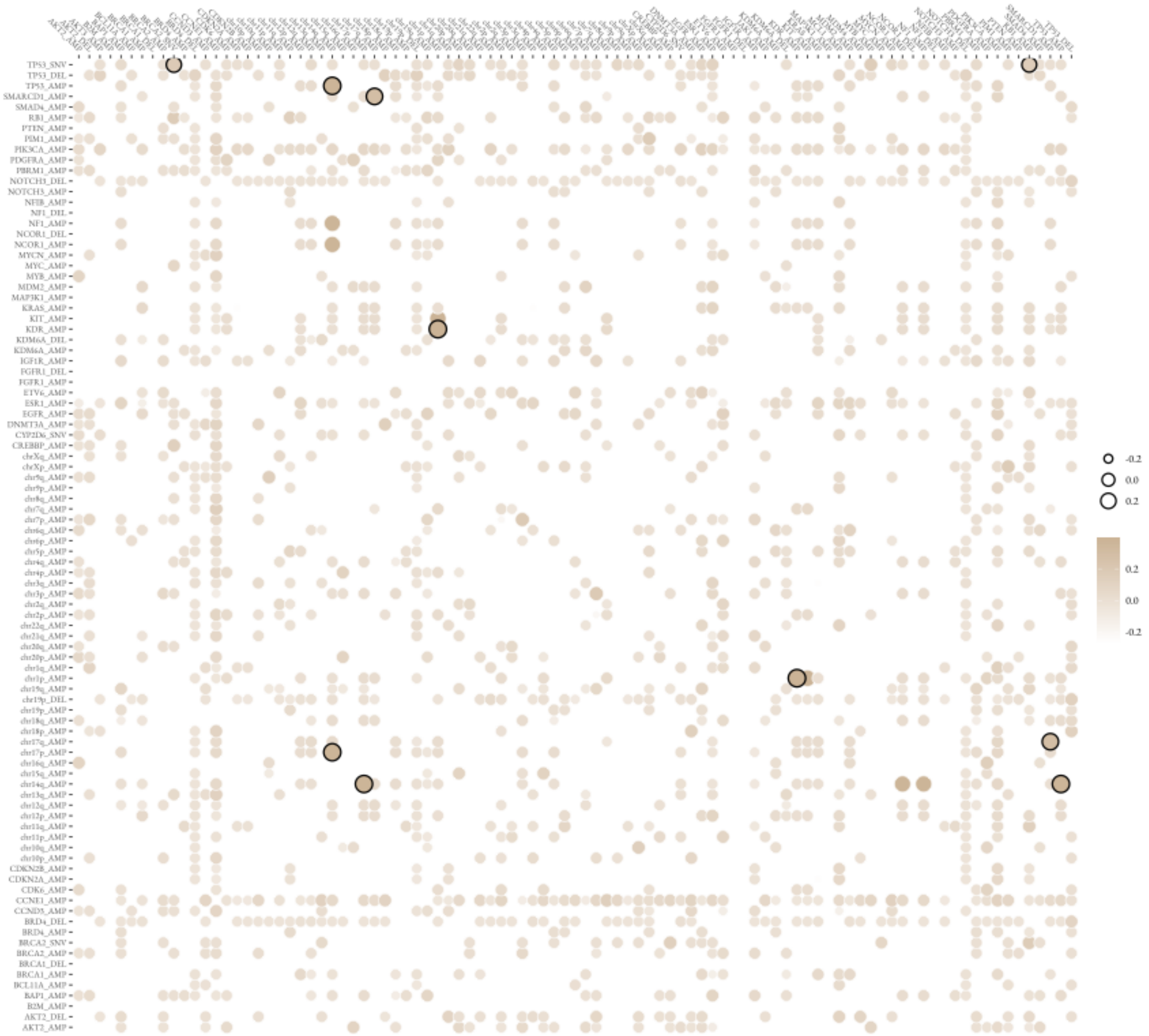


**Figure S9 | Significant co-occurrence or mutual-exclusivity of mutational events in PreT samples.** The significant interactions between chromosome arm aneuploidies, gene amplifications and SNVs. Black edges indicate a significant p-value after adjusting for false discovery rate (FDR).

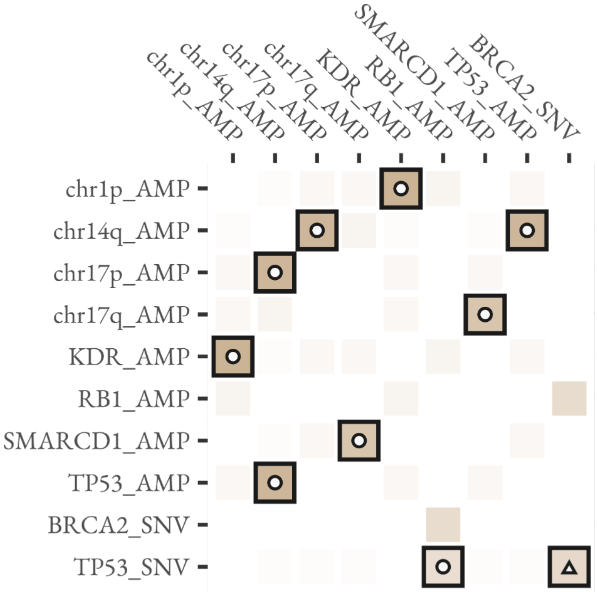


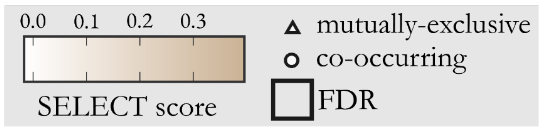


| SFE 1 | SFE 2 | Freq 1 (%) | Freq 2 (%) | Overlap (%) |
|---|---|---|---|---|
| BRCA2_SNV | TP53_SNV | 27.2 | 41.0 | 0.0 |
| RB1_AMP | TP53_SNV | 36.3 | 41 | 75.0 |
| chr14q_AMP | chr17p_AMP | 27.3 | 27.3 | 100.0 |
| chr1p_AMP | KDR_AMP | 41 | 41 | 100.0 |
| SMARCD1_AMP | chr17q_AMP | 31.8 | 31.8 | 100.0 |
| TP53_AMP | chr14q_AMP | 27.3 | 27.3 | 100.0 |

**Figure S10 | Summary of significant co-occurring or mutually-exclusive mutational events in PreT samples.** The heatmap indicates the median SELECT score of ten iterations. Interactions that are significant following FDR correction are indicated by a black outline. Circles indicate co-occurring alterations and triangles indicate alterations that are mutually exclusive. The table summarises the prevalence of each mutational event (SFE1 and SFE2) and the proportion of co-occurrence (i.e. overlap).

**Supplementary Table Legends:**

**Table S1 – Patient and Clinical characteristics**
**Table S2 – Histology Metrics**
**Table S3 – CUTseq oligos**
**Table S4 – TNBC SNV genes**
**Table S5 – SNV calls, TMB, MSI**
**Table S6 – SNV drivers**
**Table S7 – Mutational signatures**
**Table S8 – SV burden**
**Table S9 – TNBC CNA genes**
**Table S10 – GISTIC CNA**
**Table S11 – GIN, Ploidy, WGD**
**Table S12 – scarHRD results**
**Table S13 – Chromosome arm aneuploidy**
**Table S14 – LMM and GLMM Summary Table**
